# Toward a universal model for spatially structured populations

**DOI:** 10.1101/2020.12.12.422518

**Authors:** Loïc Marrec, Irene Lamberti, Anne-Florence Bitbol

## Abstract

A key question in evolution is how likely a mutant is to take over. This depends on natural selection and on stochastic fluctuations. Population spatial structure can impact mutant fixation probabilities. We introduce a model for structured populations on graphs that generalizes previous ones by making migrations independent of birth and death. We demonstrate that by tuning migration asymmetry, the star graph transitions from amplifying to suppressing natural selection. Our results are universal in the sense that they do not hinge on a modeling choice of microscopic dynamics or update rules. Instead, they depend on migration asymmetry, which can be experimentally tuned and measured.

## Introduction

Classical models of well-mixed, homogeneous microbial populations assume that each microorganism competes with all others. However, this simplification holds in few natural situations. For instance, during an infection, microbial populations are subdivided between different organs [1, 2] and hosts. Any spatial structure, e.g. that of a Petri dish, implies a stronger competition between neighbors than between distant individuals. Even well-agitated liquid suspensions feature deviations compared to idealized well-mixed populations [3].

Spatial structure can have major consequences on evolution. Remarkably, the fixation probability of a mutant can be affected, with specific structures amplifying or suppressing natural selection [4]. Studying these effects requires going beyond simple structures [5, 6] where migration is symmetric between demes (i.e. subpopulations), since fixation probabilities are unaffected in these cases [7–9], unless extinctions of demes occur [10]. Ref. [4] introduced a seminal model for complex structures, known as evolutionary dynamics on graphs, with one individual at each node of a graph, and probabilities that their offspring replaces a neighbor along each edge of the graph. However, in such models, evolutionary outcomes can drastically depend on the details of the microscopic dynamics or “update rule”, e.g. whether the individual that divides or the one that dies is chosen first, even if selection always acts at division [11–14]. This lack of universality raises issues for applicability to real populations, where one birth does not necessarily entail one death and vice-versa. Furthermore, in most microbial populations, individuals freely compete with their closest neighbors, motivating a coarse-grained description, with demes rather than individuals on graph nodes [5, 6, 15–18]. Current experiments with well-mixed demes at each node of a star graph [19] require theoretical predictions with realistic microscopic dynamics.

We propose a model for complex spatial population structures where migrations are independent from birth and death events. We investigate the fixation probability of mutants in the rare migration regime. We demonstrate that migration asymmetry determines whether the star graph amplifies or suppresses natural selection. We find a mapping to the model of Ref. [4] under specific constraints on migration rates.

### Model

We model a structured population as a directed graph where each node *i ∈* {1, …, *D*} contains a well-mixed deme with carrying capacity *K*, and migration rates *m_ij_* per individual from deme *i* to deme *j ≠ i* are specified along each edge *ij*. We then address populations including demes with different carrying capacities [20]. We consider microorganisms with two types, wild-type (W) and mutant (M), with fitnesses and death rates denoted by *f_a_* and *g_a_*, where *a* = *W* or *a* = *M*. Here, we call fitness the maximal division rate of microorganisms, reached in exponential growth. Their division rate in deme *i* is given by the logistic function *f_α_*(1 − *N_i_*/*K*), where *N_i_* is the number of individuals in deme *i*. We take wild-type fitness as a reference, *f_W_* = 1. We address selection on birth, and hence *g_M_* = *g_W_*, but our results can be generalized to selection on death. We focus on the regime where deme sizes *N_i_* fluctuate weakly around their deterministic steady-state values, without extinctions [10, 21, 22].

We assume that mutations are rare enough for further mutation events to be neglected while the fate of a given mutant lineage (taking over or disappearing) is determined. We consider an initial mutant placed uniformly at random, which is realistic for spontaneous mutations occurring either with a fixed rate or with a fixed probability upon division. Note that in models with one individual per node, uniform initialization is more appropriate in the first case, while placing mutants proportionally to the replacement probability of a node (“temperature initialization”) is more appropriate in the second one [23]. This distinction vanishes here, as division rate does not depend on location. Under uniform initialization, the fixation probability of a neutral mutant is independent of structure for connected graphs [20]. Compared to the well-mixed population with the same total size, an *amplifier* of natural selection features a larger fixation probability for beneficial mutants (*f_M_* > *f_W_*), and a smaller one for deleterious mutants (*f_M_* < *f_W_*), while a *suppressor* has the opposite characteristics [24].

We focus on the rare migration regime [9], where fixation of a type (W or M) in a deme is much faster than migration timescales. Then, the state of the population can be described in a coarse-grained way by whether each deme is mutant or wild-type. Its evolution is a Markov process where elementary steps are migration events, which change the state of the system if fixation ensues. Then, a mutant first needs to fix in the deme where it appeared before spreading. Since fixation in a homogeneous deme is well-known, we study the second stage, starting from one fully mutant deme.

### Link with models with one individual per node

A formal mapping can be made between our model and that of [4], if the same graph is considered, with a deme per node in our model and with one individual per node in [4] (see [20]). The probability 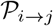 that, upon a migration event resulting into fixation, an individual from deme *i* takes over in deme *j* in our model maps to the probability 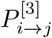 that, upon a division, the offspring from node *i* replaces the individual on node *j* in the model of [4]:

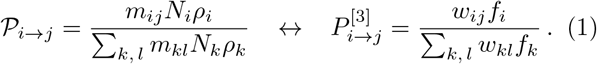

In this mapping, the product 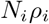 of deme size *N_i_* and fixation probability *ρ_i_* of an organism from deme *i* in our model plays the part of fitness *f_i_* of the individual on node *i* in [4], while the migration rate *m_ij_* plays the part of the replacement probability *w_ij_* that the offspring of the individual in *i* replaces that in *j*. However, an important constraint in the “Birth-death” model of [4] (also known as biased invasion process [11]) is Σ_*j*_ *w_ij_* = 1 for all *i*, because replacement includes birth, migration and death at once, and population size is constant. By contrast, migration rates *m_ij_* in our model are all independent.

A generalized circulation theorem holds for our model [20], in the spirit of [4]. Specifically, a population of *D* demes on a graph has the same mutant fixation probability as the clique if and only if, for all nodes of the graph, the total outgoing migration rate is equal to the total incoming migration rate.

Thus, we expect fixation probabilities in our model to map to those of [4] for circulations or if Σ_*j*_ *m_ij_* is independent of *i*, but to potentially differ otherwise. We now consider specific graphs with strong symmetries.

### Clique and cycle

In the clique (or island model [5, 6]), all demes are equivalent and connected to all others with identical migration rates *m* per individual (Fig. 1, upper inset). Starting from one fully mutant deme and *D* − 1 fully wild-type demes, the fixation probability 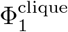 of the mutant reads [20] (proof inspired by [9, 25]):

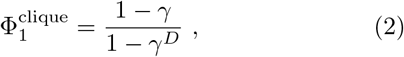

with

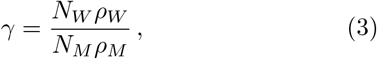

where *N_W_* (resp. *N_M_*) is the deterministic steady-state size of a wild-type (resp. mutant) deme and *ρ_W_* (resp. *ρ_M_*) is the fixation probability of a wild-type (resp. mutant) microbe in a mutant (resp. wild-type) deme. This result is independent of migration rate *m*, and Eq. (2) has the exact same form as the fixation probability of a single mutant in a well-mixed population of fixed size *D* in the Moran model [26, 27], but with *γ* playing the role of the ratio *f_W_* / *f_M_*, consistently with the formal mapping Eq. (1) between our model and that of [4] where *N_ρ_* plays the part of fitness. 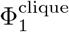 is plotted versus *f_M_* in Fig. 1, showing excellent agreement between Eq. (2) and our stochastic simulation results. Moreover, this fixation probability is very close to that in a well-mixed population. We show [20] that the clique is a slight suppressor of selection, but that modeling migrations as exchanges of individuals and assuming *N_M_* = *N_W_* exactly recovers the well-mixed result, consistently with results on symmetric migrations [7, 8].

**FIG. 1.**
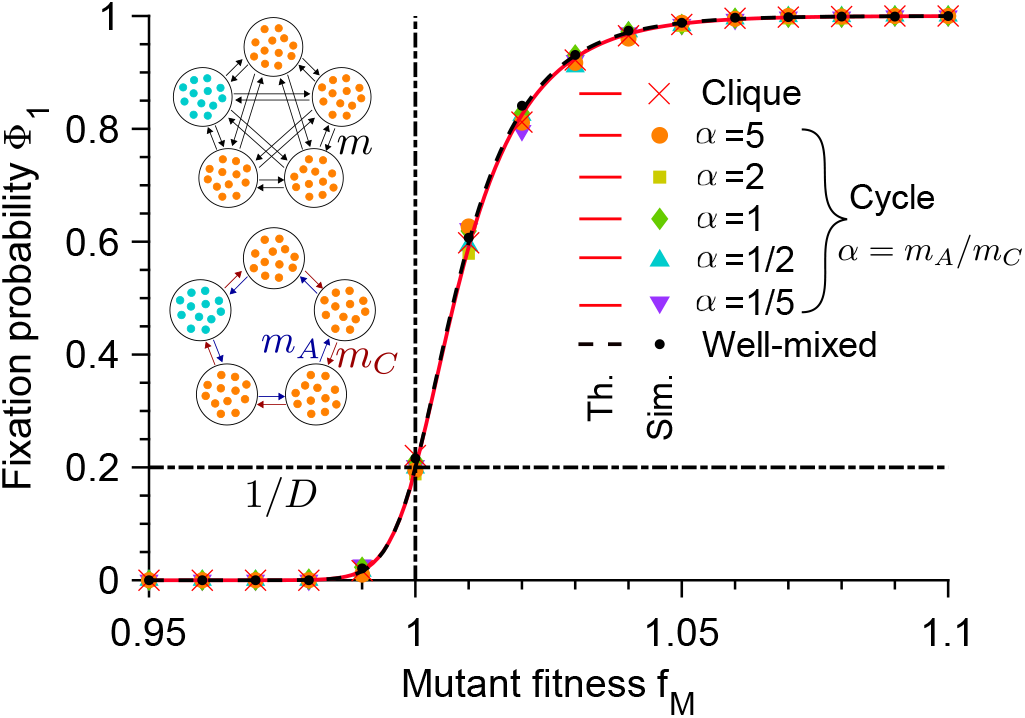
Fixation probability Φ_1_ of the mutant type versus mutant fitness *f_M_*, for the clique (see upper inset), and for the cycle (see lower inset) with different migration rate asymmetries *α* = *m_A_*/*m_C_*, starting with one fully mutant deme. Data for the well-mixed population is shown as reference, with same total population size and initial number of mutants. Markers are computed over 10^3^ stochastic simulation realizations. Curves represent analytical predictions, Eq. (2) for the cycle and the clique, and Eq. (S15) [20] for the well-mixed population [26, 27]. Vertical dash-dotted lines indicate the neutral case *f_M_* = *f_W_*, and horizontal dash-dotted lines represent the neutral fixation probability. Parameter values: *D* = 5, *K* = 100, *f_W_* = 1, *g_W_* = *g_M_* = 0.1 in both panels. In simulations, for the clique, *m* = 10^−6^; for the cycle, from top to bottom, (*m_A_*, *m_C_*) × 10^6^ = (1, 5); (1, 2); (1, 1); (2, 1); (5, 1).

Another graph where all demes are equivalent is the cycle. Clockwise and anti-clockwise migrations can have different rates, denoted respectively by *m_C_* and *m_A_* (Fig. 1, lower inset). The cycle resembles the circular stepping-stone model [7], but can feature asymmetric migrations. We show [20] that the fixation probability 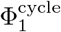 is the same as for the clique, Eq. (2), as corroborated by our simulations, see Fig. 1. Indeed, the cycle is a circulation. In particular, migration rates do not impact 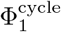.

### Star

In the star, a central node is connected to all others, called leaves. An individual can migrate from a leaf to the center with migration rate *m_I_* and vice-versa with rate *m_O_* (Fig. 2, inset). The mutant fixation probability can be expressed exactly as a function of *D*, *α* = *m_I_*/*m_O_* and *γ* defined in Eq. (3) (proof [20] inspired by [28]):

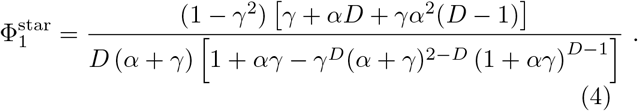

**FIG. 2.**
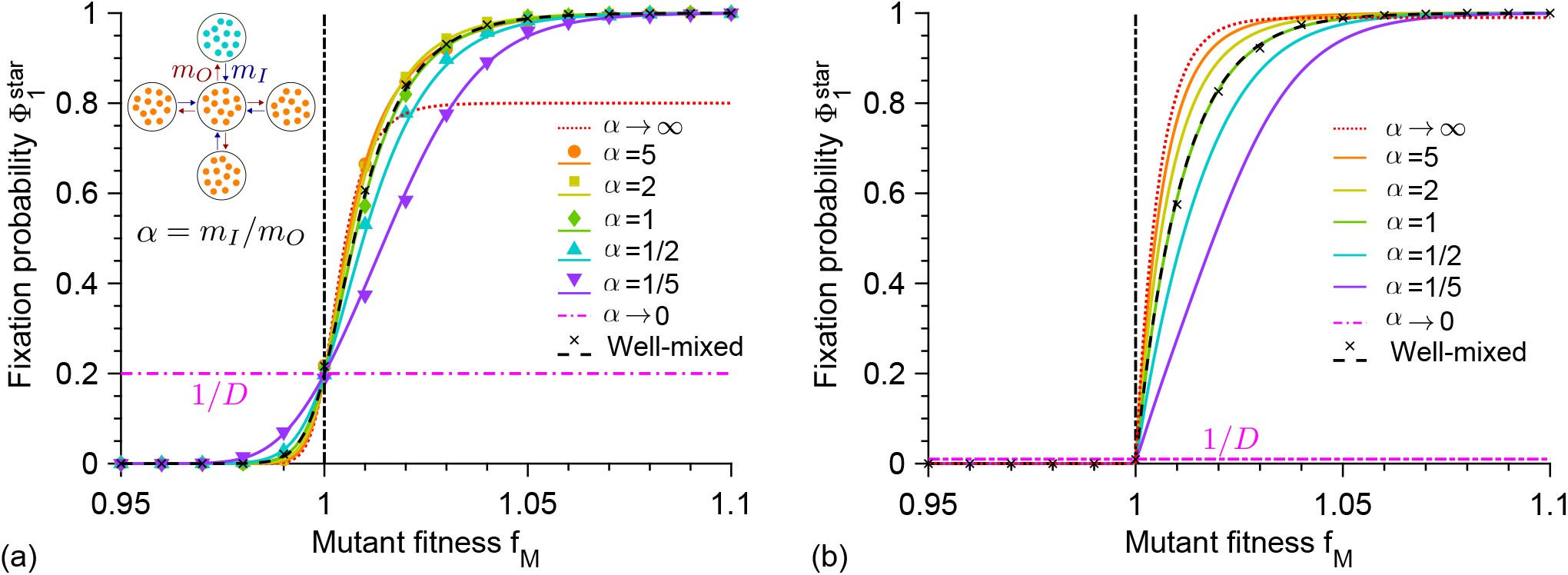
Fixation probability 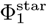 of the mutant type in a star graph versus mutant fitness *f_M_*, starting with one fully mutant deme chosen uniformly at random, for different migration rate asymmetries *α* = *m_I_*/*m_O_*. Number of demes: *D* = 5 (a) and *D* = 100 (b). Data for the well-mixed population is shown as reference, with same total population size and initial number of mutants. Markers are computed over 2 × 10^3^ stochastic simulation realizations. Curves represent analytical predictions in Eq. (4). Vertical dash-dotted lines indicate the neutral case *f_W_* = *f_M_*. Parameter values: *K* = 100, *f_W_* = 1, *g_W_* = *g_M_* = 0.1. In panel (a), from top to bottom, (*m_I_*, *m_O_*) × 10^6^ = (5, 1); (2, 1); (1, 1); (1, 2); (1, 5) in simulations.

Fig. 2(a) shows the fixation probability 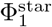 of the mutant type for different values of migration asymmetry *α* = *m_I_*/*m_O_*, with very good agreement between Eq. (4) and our simulations. If *α* < 1, the star suppresses selection compared to the clique, while for *α* > 1 it slightly amplifies selection in some range of mutant fitness *f_M_* [20]. For *α* = 1, 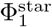 reduces to the fixation probability of the clique, Eq. (2) [20]. Consistently, the star is a circulation for *α* = 1. Stronger amplification for *α* > 1 is obtained for large *D* (Fig. 2(b)). Qualitatively, for large *D*, mutants very likely start in a leaf. If *α* is large, they often spread to the center, which helps fit mutants take over. Conversely, if *α* is small, the center often invades the leaves, thus preventing any mutant originating in a leaf from fixing. Results with mutants starting in a specific deme are also shown in [20].

Imposing that Σ_*j*_ *m_ij_* is independent of *i* amounts to imposing *α* = *D* − 1 in the star [20]. Then, Eq. (4) reduces to that [28] obtained in the model of [4], with *γ* in Eq. (3) playing the role of *f_W_*/*f_M_* [20], as per our general mapping Eq. (1). The celebrated amplification property of the star in the large *D* limit [4, 29] is thus exactly recovered in our model for *α* = *D* − 1.

While the star is an amplifier for large *D* in the model of [4], it can either suppress or an amplify selection, depending on *α*, in our model where *D* and *α* are two independent parameters. Fig. 3 shows that restricting to *α* = *D* − 1 yields amplification. In models with one individual per node, the star is an amplifier for large *D* for the Birth-death dynamics (“update rule”), where one individual is chosen to divide and its offspring replaces one of its neighbors [4], but a suppressor for the death-Birth dynamics (or biased voter model [11]), where one individual is chosen to die and is replaced by the offspring of one of its neighbors (selection being on division rates in both cases, as denoted by the upper-case “Birth” [13], and resulting in global selection in the Birth-death case and local selection in the death-Birth case) [13, 24, 30, 31]. Consistently, the latter dynamics would yield *α* = 1/(*D* − 1).

**FIG. 3.**
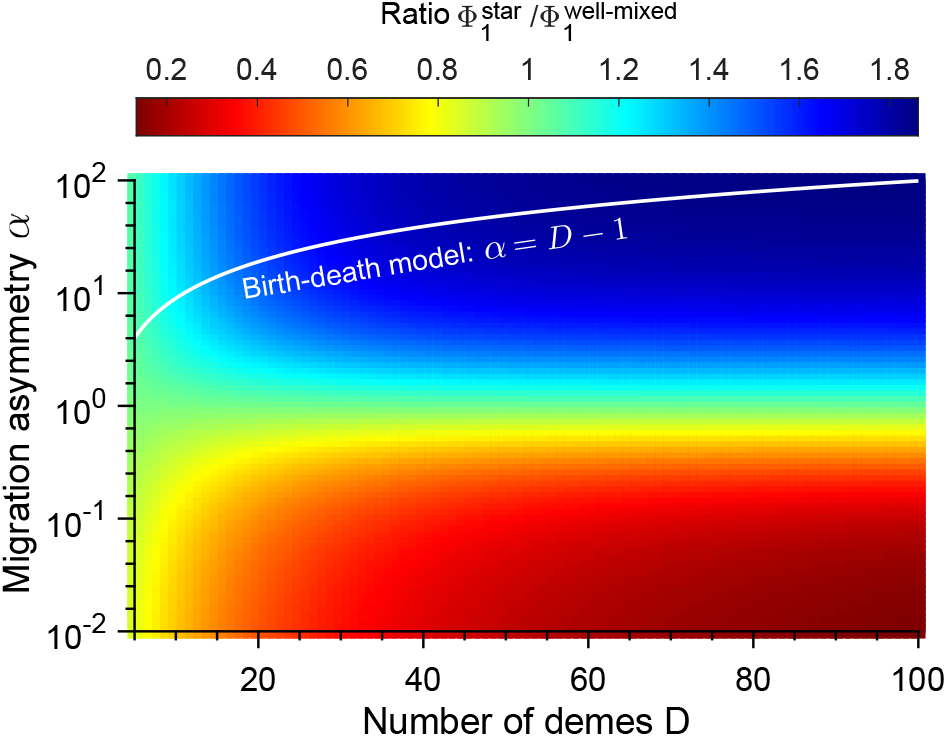
Amplification and suppression properties for the star. Heatmap of the ratio of the fixation probability 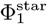 of the mutant type in a star graph to that 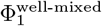 in a well-mixed population with same total population size and initial number of mutants, versus number *D* of demes and migration rate asymmetry *α* = *m_I_*/*m_O_*. The star is initialized with one fully mutant deme chosen uniformly at random. Data from analytical formula in Eq. (4) for the star, and in Eq. (S15) [20] for the well-mixed population. Parameter values: *K* = 100, *f_W_* = 1, *f_M_* = 1.001, *g_W_* = *g_M_* = 0.1.

### Comparison to [16]

A model generalizing [4] to graphs where each node contains a deme with fixed population size was introduced in [16] (see also [15, 18, 32]). In this model, as in [4], each elementary event comprises a birth in one deme and a death in another one, yielding Birth-death and death-Birth models that give different results. Rare migrations in our model correspond to strong self-loops (migrations to the original deme) in the model of [16]. For the star [23], we show [20] that by matching migration-to-division rate ratios in each deme, both models yield similar simulation results. However, even then, a difference is that death rate (resp. birth rate) is not homogeneous across demes in the Birth-death (resp. death-Birth) models of [16], unless migrations are symmetric. Our model allows more realistic choices.

## Discussion

We developed a model of spatially structured microbial populations on graphs where migration, birth and death are independent events. We showed that for rare migrations, the star graph continuously transitions between amplifying and suppressing natural selection as migration rate asymmetry is varied. This elucidates the apparent paradox in existing models, where the star, like many random graphs [13], is an amplifier in the Birth-death dynamics and a suppressor in the death-Birth dynamics [13, 24, 30, 31]. We found a mapping between our model and that of [4], under a constraint on migration rates. Models with one individual per node require making specific choices on the microscopic dynamics (“update rule”), which constrain migration rates. By lifting this constraint, our model reconciles and generalizes previous results, showing that migration rate asymmetry is key to whether a given population structure amplifies or suppresses natural selection. This crucial role of migration asymmetry is consistent with the fact that structures with symmetric migrations do not affect fixation probabilities [7, 8].

Birth-death dynamics may be realistic for extreme resource limitation, such that one birth causes one death, while death-Birth dynamics may better model cases where death frees resources, e.g. light for plants [30, 33]. However, in general, in a microbial population, population size is not strictly fixed, and the order of birth and death events is not set. In our more universal model, results do not hinge on a modeling choice made for microscopic dynamics. Instead, they depend on a quantity that can be directly set or measured in experiments, namely migration rate asymmetry. The differences between Birth-death and death-Birth dynamics are major for mutant fixation probabilities, but also in evolutionary game theory, where spatial structure can promote the evolution of cooperation in the latter case, but not in the former [34–36]. Previous efforts were made to generalize beyond these dynamics by allowing both types of update to occur in given proportions [37, 38]. Interestingly, it was recently shown that no general amplification of selection can occur when even a small proportion of death-Birth events occurs [38], in contrast to the Birth-death case. Conversely, in our model, the amplification property of the star graph in the large-size limit is preserved, but for sufficient migration asymmetry.

While our focus was on mutant fixation probabilities, our model can be employed to investigate fixation times and evolution rate [18, 39–45]. It can also address more complex population structures [24, 46], e.g. motivated by within-host or between-host pathogen dynamics [47]. Our study can be extended beyond the regime of rare migrations [48], and to models of evolutionary game theory, as well as to diploid organisms [10, 21, 22, 49, 50]. Finally, our work allows direct comparisons with quantitative experiments [19]. Other experiments could be performed using e.g. microfluidic devices allowing to control flow between different populations [51], or microtiter plates where dilutions and migrations can be performed via a liquid-handling robot [52–54]. Applications in biotechnology could be envisioned, e.g. amplifying *in vivo* selection in the directed evolution of biomolecules [55].

## Acknowledgments

This project has received funding from the European Research Council (ERC) under the European Union’s Horizon 2020 research and innovation programme (grant agreement No. 851173, to AFB). LM acknowledges funding by a graduate fellowship from EDPIF. LM thanks his grandfather, Jean Polard, for inspiration.

## I. FIXATION PROBABILITY OF NEUTRAL MUTANTS

Consider a graph made of nodes *i*, each associated to a deme with steady-state population size *N_i_*, and edges *ij* where migration rates *m_ij_* from deme *i* to deme *j* are specified. Further assume that the graph is not disconnected. Consider uniform initial conditions: a mutant has probability *N_i_*/Σ_*j*_ *N_j_* to be initially placed in deme *i*. A neutral mutant then has probability 1/*N_i_* to fix in deme *i* (taking the result for constant population size [27]). Let 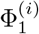 denote the probability that the mutant fixes in the whole metapopulation, starting from a fully mutant deme *i*, all other demes being fully wild-type. The overall fixation probability of one mutant in the metapopulation reads

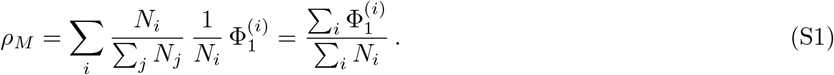

Let us now remark that, because the graph is not disconnected, after a sufficient time, all individuals in the metapopulation are descended from the same deme. If we start from one deme *i* that is fully mutant and all others that are fully wild-type, this yields

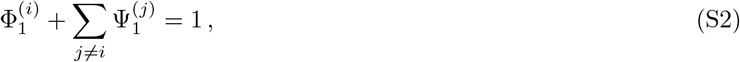

where 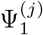 is the probability that wild-type individuals from deme *j* fix in the whole metapopulation. But since the mutant is assumed to be neutral, we have 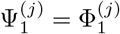, and thus Eq. S2 becomes

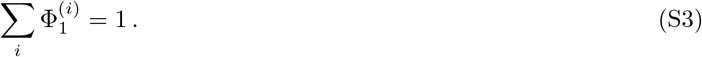

Therefore, combining Eqs. S1 and S3, we obtain

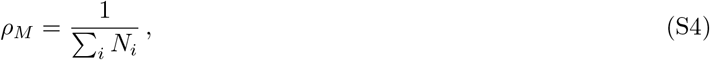

which is exactly the fixation probability of one neutral mutant in a well-mixed population of size Σ_*i*_ *N_i_* [27]. Thus, provided that the graph is not disconnected, the fixation probability of a neutral mutant under uniform initial conditions is independent of population structure in our model.

In the particular case where all *D* demes have the same size, i.e. *N_i_* ≡ *N* does not depend on *i*, then *ρ_M_* = 1/(*ND*), and the average fixation probability starting from one single fully mutant deme under uniform initial conditions is

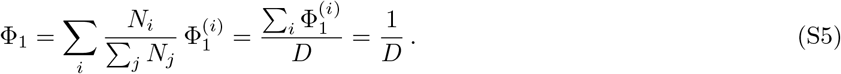

## II. FIXATION PROBABILITIES IN STRONGLY SYMMETRIC GRAPHS

In this study, we investigate the fate of mutants in the structures shown in Fig S1. The clique, cycle and star are considered in the present section, while the doublet, shown in panel D, is tackled in section VI as it involves demes with different population sizes.

**FIG. S1.**
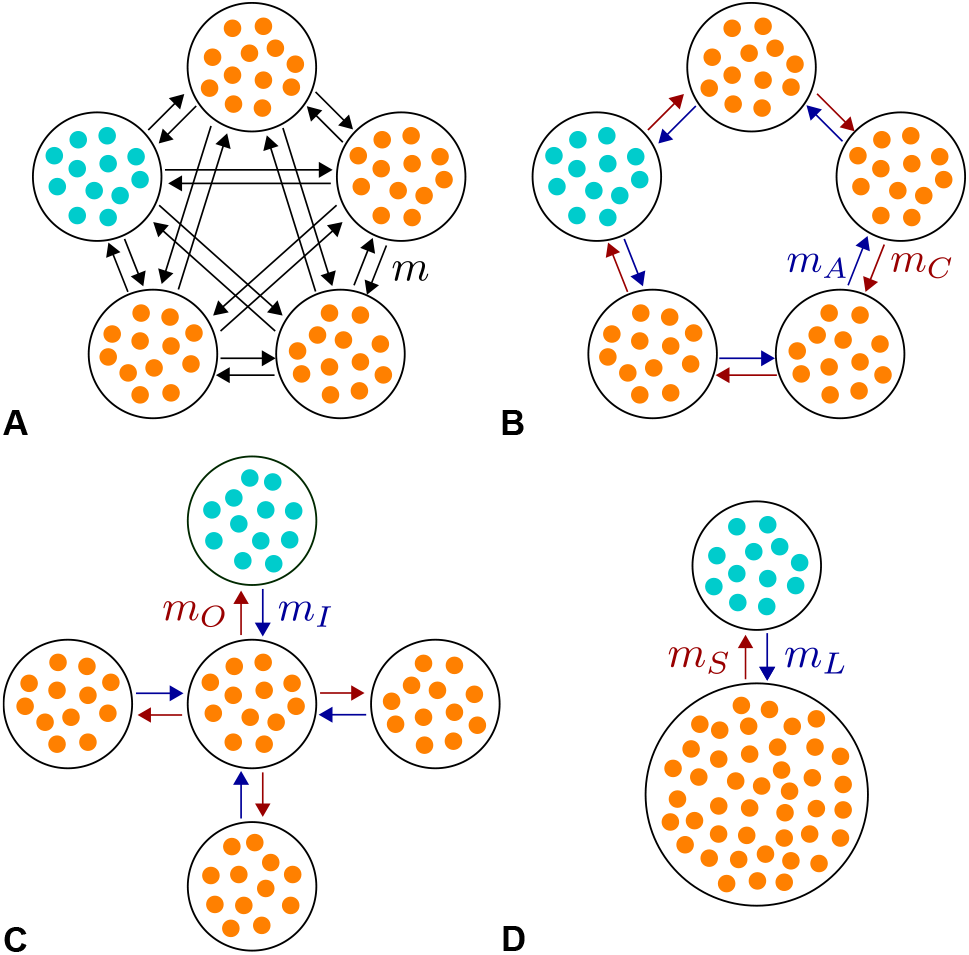
Some population structures. **A:** Clique. **B:** Cycle. **C:** Star. **D:** Doublet comprising a small deme and a larger deme. Mutants (M) are in blue, wild-type (W) in orange, and the state where the mutant type has fixed in one deme while all other demes are fully wild-type is represented. Arrows indicate migrations, with the associated rates per individual.

### A. Clique

#### 1. General expression

Let us consider a population with *D* demes, structured as a clique, i.e. where migration rates per individual between all demes are equally likely (Fig. S1A). The state of the system can be fully described by the number *i* of mutant demes. We denote by *m* the migration rate per individual from one deme to any other deme. Recall that in our model, migration occur between two different demes (no migration can end in the deme where it started). Let us assume that we start from *i* fully mutant demes and *D* − *i* fully wild-type demes. Recall that the wild-type is denoted by *W* and the mutant by *M*.

Consider the outcome of a migration event. The number of mutant demes increases by 1 if an *M* individual migrates from one of the *i* mutant demes to one of the *D − i* wild-type demes, and fixes there. The probability that this occurs upon a migration event thus reads

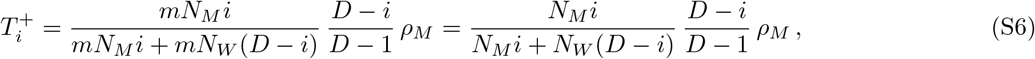

where

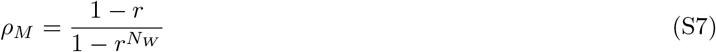

is the fixation probability of a mutant microbe in a wild-type deme in the Moran process [26, 27] (see section VII) where

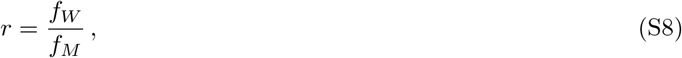

and *N_W_* = *K*(1 − *g_W_*/*f_W_*) is the steady-state size of a wild-type deme. Similarly, the number of mutant demes decreases by 1 if a *W* individual migrates from one of the *D* − *i* wild-type demes to one of the *i* mutant demes, and fixes there. The probability that this occurs upon a migration event thus reads

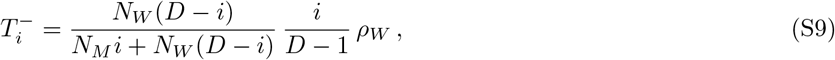

where

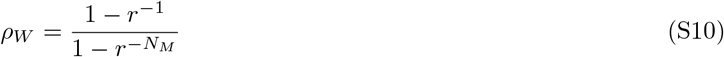

is the fixation probability of a wild-type microbe in a mutant deme, with *r* in Eq. S8, and *N_M_* = *K*(1 − *g_M_*/*f_M_*) is the steady-state size of a mutant deme.

The fixation probability 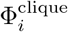 of the mutant type in a clique of *D* demes starting with *i* fully mutant demes satisfies the recurrence relation

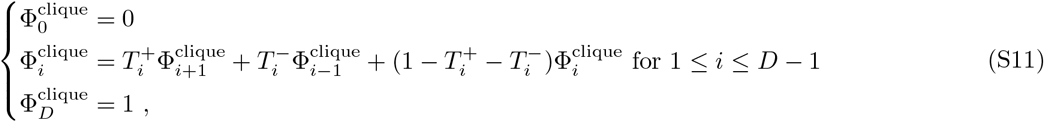

where the second equation follows from distinguishing the different outcomes of the first migration event. Eq. S11 can be solved e.g. as in Ref. [25], yielding

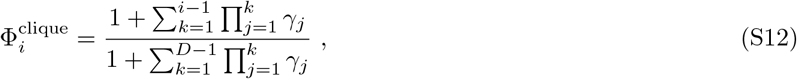

where 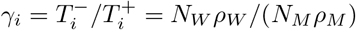. Since here *γ_i_* does not depend on the initial number *i* of mutant demes, the fixation probability 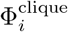 reduces to

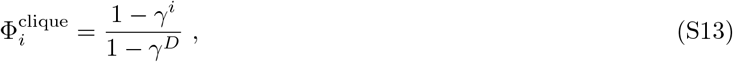

with

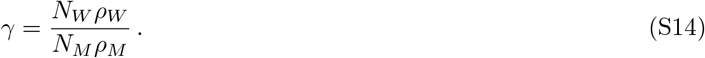

Note that in the neutral case where *γ* = 1, Eq. S12 yields 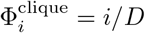, consistently with Eq. S5.

Eq. S13 has the same form as the fixation probability of *i* mutants in a well-mixed population of *N* individuals in the Moran model [26, 27], namely

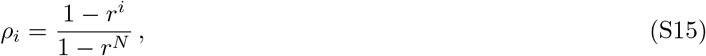

with *r* in Eq. S8 (note that Eq. S7 corresponds to the case *i* = 1). Specifically, Eq. S15 maps to Eq. S13 by replacing *N* with *D* and *r* with *γ*. Thus, the clique can be thought of a coarse-grained version of the well-mixed population (see also Ref. [9]): each deme is identically connected to all other demes, just like all individuals are in competition in the well-mixed population.

To obtain the fixation probability 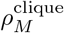 of a single mutant individual in the clique, one needs to include the fixation of the mutant in one deme before the spread from that deme to the full metapopulation. Thus, it reads

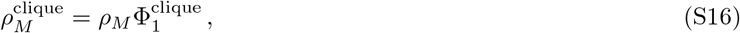

with *ρ_M_* given by Eq. S7 and 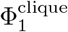 by Eq. S13 for *i* = 1.

#### 2. Expansion for very small mutational effects

Let *ϵ* be such that *f_M_* = *f_W_* (1 + *ϵ*). Consider the regime where *ϵ* ≪ 1 and *N_W_*|*ϵ*| ≪ 1. Then Eq. S13 gives (for *i* = 1)

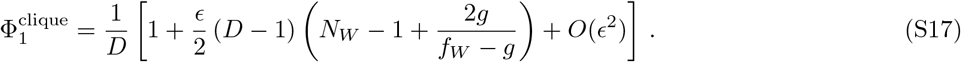

Eqs. S15 and S16 then allow us to show that

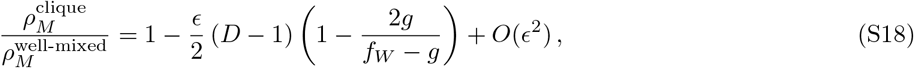

where 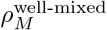 is the fixation probability of a mutant in a well-mixed population with *N_W_ D* individuals. Therefore, the clique is a suppressor of natural selection in this regime if *g* < *f*/3. Suppression is all the more important that the degree of subdivision is high, namely the number *D* of demes (recall that here we compare a clique and a well-mixed population for the same total population size *N_W_ D*).

#### 3. Expansion for relatively small mutational effects

Next, consider the regime where *ϵ* ≪ 1 and *N_W_* |*ϵ*| ≫ 1 but *N_W_*ϵ*^2^* ≪ 1. Then, if *ϵ* > 0,

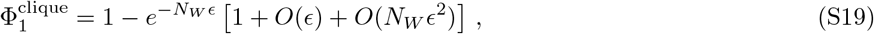

which means that fixation is almost certain (recall that if one starts from one single mutant, this holds provided that a mutant has fixed in a deme, which occurs with probability *ρ_M_* = *ϵ* + *O*(*ϵ*^2^), see Eq. S7). Therefore, Eq. S16 yields 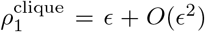, which is equal (to this order) to the fixation probability 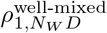 in a well-mixed population with the same total size *N_W_ D*. Now, if *ϵ* < 0,

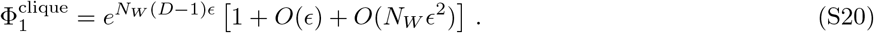

which means that fixation is exponentially suppressed. Thus, Eq. S16 yields 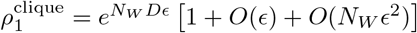, which is equal (to this order) to the fixation probability 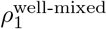 in a well-mixed population with the same total size *N_W_ D*. Hence, in this regime, the fixation probability of a mutant in the clique is very close to that in a well-mixed population with the same total size *N_W_ D*, and the suppression effect found for extremely small mutational effects is quite restricted.

#### 4. Model specialized to symmetric migrations

In the clique, migrations are symmetric, i.e. *m_ij_* = *m_ji_* = *m* for all *i* ≠ *j*. Let us consider another model, restricted to symmetric migrations, where each migration event is modeled as an exchange between two individuals from two different demes. Let us further neglect the difference between *N_M_* and *N_W_*, and assume *N_M_* = *N_W_* = *N*. Upon a given migration event, the probability that the number *i* of mutant demes increases is

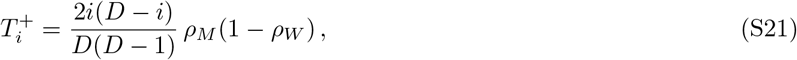

and similarly, the probability that *i* decreases is

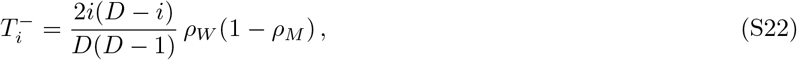

yielding as above the fixation probability 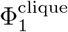 in Eq. S13 when one starts from one mutant deme, but with

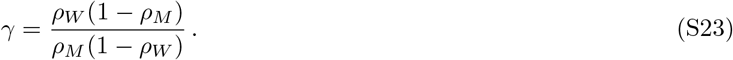

Eq. S16 then yields the fixation probability of one single mutant

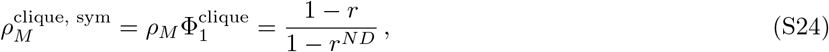

with *r* defined in Eq. S8. This is exactly the fixation probability of a mutant in a well-mixed population of fixed size *ND* in the Moran model (see above).

### B. Cycle

Let us consider a population structured as a cycle with *D* demes (see Fig. S1B), starting from exactly one fully mutant deme. During the fixation process, this will yield a cluster of *i* consecutive mutant demes that cannot break. Therefore, in this process, the state of the system can be fully described by the number *i* of (consecutive) mutant demes. Upon a migration event, the number *i* of mutant demes increases by 1 if a *M* individual from one of the two extremities of the mutant cluster migrates to the neighboring wild-type deme and fixes there. The probability that this occurs thus reads

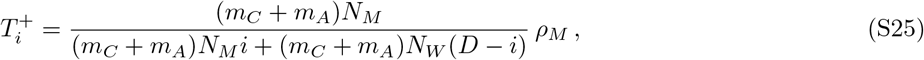

with *ρ_M_* given by Eq. S7. Similarly, the number *i* of mutant demes decreases by 1 if a *W* individual from either of the two wild-type demes surrounding the mutant cluster migrates and fixes in its neighboring mutant deme. The probability that this occurs upon a migration event thus reads

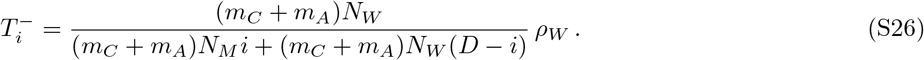

Thus, the fixation probability Φ_*i*_ of mutation starting with *i* consecutive mutant demes satisfies Eq. S11 with 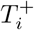 and 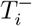 given by Eqs. S25 and S26, which yields the fixation probability in Eq. S13 with *γ* given by Eq. S14. The fixation probability in the cycle, starting from exactly one fully mutant deme, is thus equal to that of the clique with the same number of demes.

### C. Star

#### 1. General expression

Let us consider a population structured as a star with *D* demes (see Fig. S1C). Migrations from each single leaf to the center occur with a rate per individual *m_I_* while migrations from the center to each single leaf occur with a migration rate per individual *m_O_*. The state of the system can be fully described by a binary number indicating whether the center is wild-type or mutant and the number *i* of mutant leaves.

Upon a given migration event, the probability that the mutant type fixes in the center, if the center is initially wild-type and *i* leaves are mutant, reads

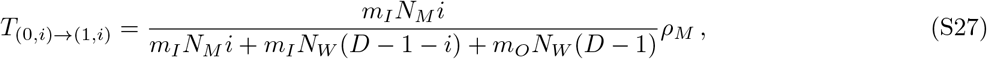

because it happens if migration occurs from a mutant leaf to the center, and the mutant then fixes in the center. Similarly, the probability that the wild-type fixes in the center, if the center is initially mutant and *i* leaves are mutant, reads

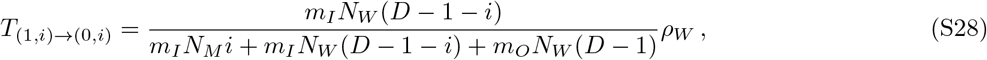

while the probability that the number of mutant leaves increases by 1 if the center is mutant is

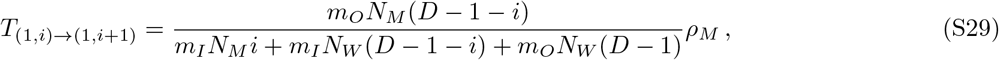

and the probability that the number of mutant leaves decreases by 1 if the center is wild-type is

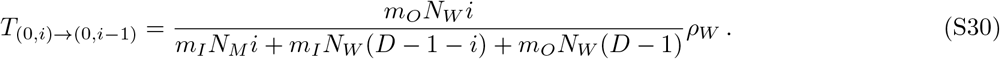

Let 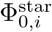 be the fixation probability of the mutant type starting from *i* fully mutant leaves and a wild-type center. Similarly, let 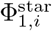 be the fixation probability of the mutant type starting from *i* fully mutant leaves and a mutant center. The fixation probabilities 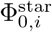 and 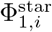 satisfy the following recurrence relationship, which is analogous to that in Ref. [28]:

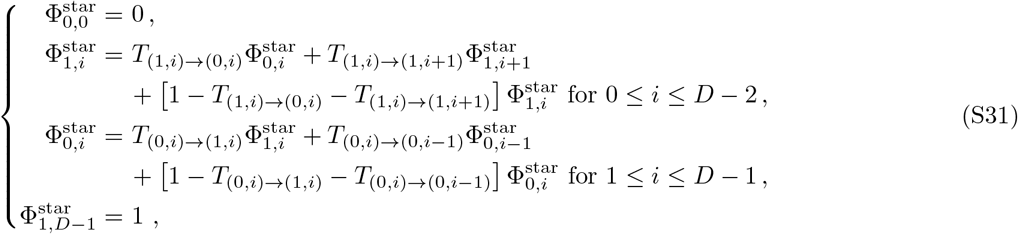

Employing the expressions of the transition probabilities given above, the system S31 can be rewritten as:

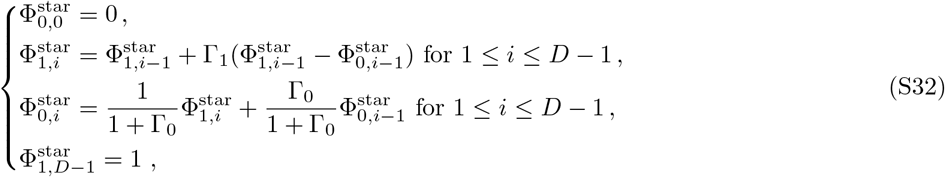

where Γ_1_ = *γ m_I_*/*m_O_* and Γ_0_ = *γ m_O_*/*m_I_*, with *γ* given in Eq. S14. Solving the system S32 yields

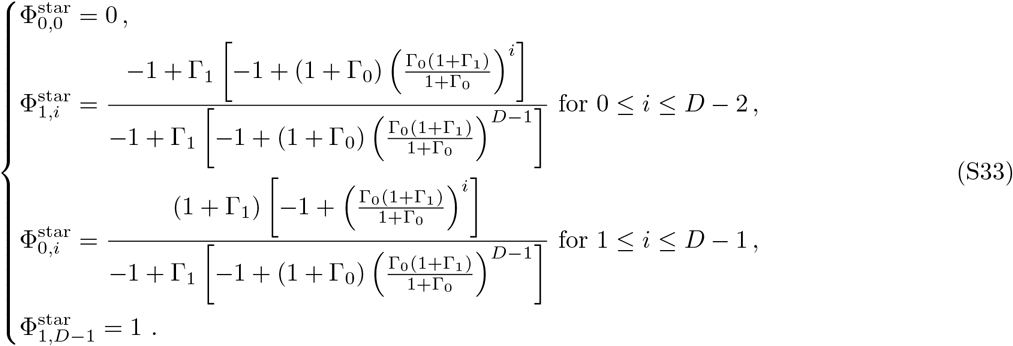

In particular, the fixation probability of the mutant type starting from one fully mutant center and all leaves fully wild-type reads

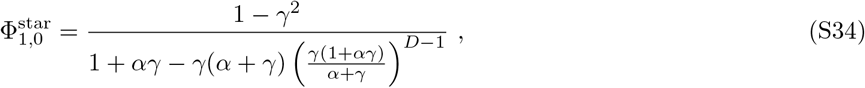

where

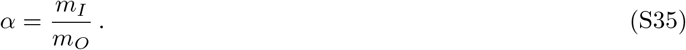

The fixation probability of the mutant type starting from one fully mutant leaf and all other demes fully wild-type reads

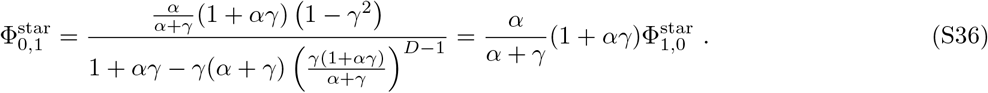

The probability that the mutant type fixes, starting from a mutant deme that can be any deme of the star with equal probability, can then be expressed as

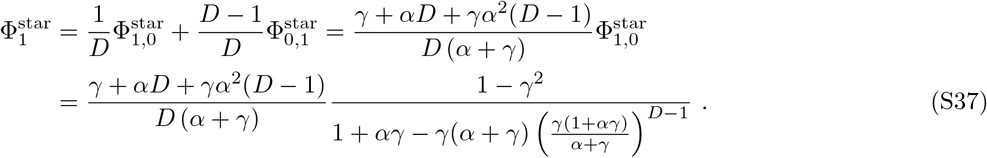

which can be rewritten as Eq. 4 in the main text.

Importantly, Eqs. S34, S36 and S37 are exactly equivalent to the formula given in Ref. [28] if the following substitutions are made: *γ* → *f_W_*/*f_M_* and *α* → *D* − 1 (bearing in mind that in the notations of Ref. [28], *D* − 1 is called *n* and *f_W_*/*f_M_* is called 1/*r*).

#### 2. Expansion for very small mutational effects

Let *ϵ* be such that *f_M_* = *f_W_* (1 + *ϵ*). For uniform initialization, consider the regime where *ϵ* ≪ 1 and *N_W_* |*ϵ*| ≪ 1. Then Eq. S37 yields

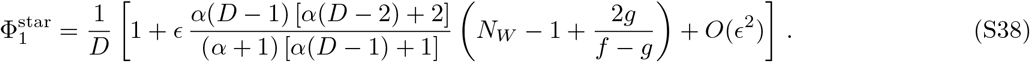

Comparing Eqs. S17 and S38 yields

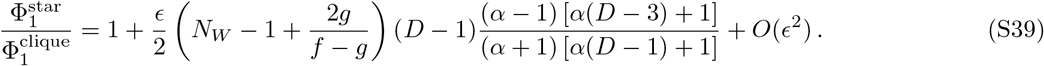

Assuming *D* > 2, the first-order term in Eq. S39 has the same sign as *ϵ*(*α* − 1). Thus, in this regime, the star is an amplifier of selection with respect to the clique for *α* > 1, and a suppressor for *α* < 1. Furthermore, for *D* > 3 (and integer), the function

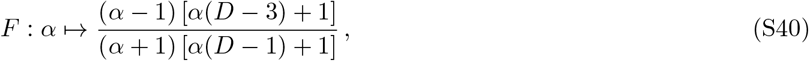

increases with *α* for *α* > 0, which entails that, for very small mutational effects, the strongest amplification is obtained for *α* ≫ 1, where *F* (*α*) → 1. Conversely, if *α* ≪ 1 and *α* ≪ 1/*D*, Eq. S38 yields

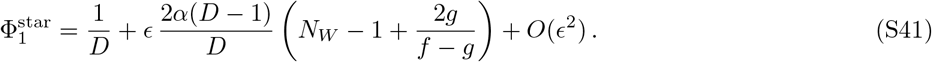

In particular, the coefficient of the first-order term in *ϵ* becomes very small if *α* ≪ 1/ (2*N_W_*), meaning that for such small values of *α*, we expect a strong suppression of selection, with a fixation probability that becomes independent of *ϵ* and flat (for very small mutational effects *ϵ*).

#### 3. Expansion for relatively small mutational effects

Next, consider the regime where *ϵ* ≪ 1 and *N_W_* |*ϵ*| ≫ 1 but *N_W_*ϵ*^2^* ≪ 1. Then, if *ϵ* > 0, Eq. S37 yields

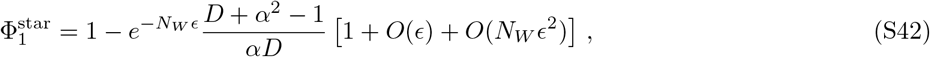

which gives, employing Eq. S19,

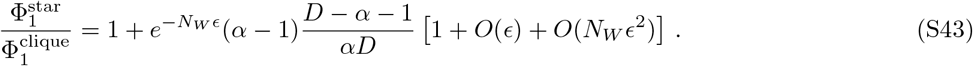

Thus, in this case, assuming *D* > 2, we have 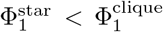 if *α* < 1 or *α* > *D* − 1, whereas 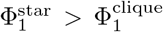 if 1 < *α* < *D* − 1. Now if *ϵ* < 0, Eq. S37 yields

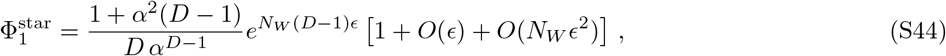

which gives, employing Eq. S20,

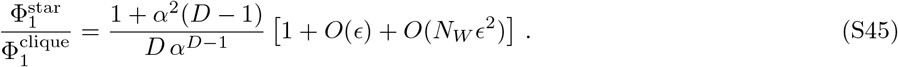

Then, assuming *D* > 3, we have 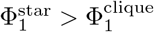 if *α* < 1 while 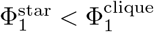 if *α* > 1.

Combining results for *ϵ* > 0 and *ϵ* < 0 in this regime, as well as results obtained for very small mutational effects above, we find that the star is a suppressor of selection compared to the clique for *α* < 1, an amplifier of selection for 1 < *α* < *D* − 1, and a transient amplifier of selection for *α* > *D* − 1. Indeed, in the latter case, one switches from amplification to suppression as *ϵ* is increased. Specifically, there is amplification for *ϵ* < 0 satisfying *ϵ* ≪ 1 and *N_W_* |*ϵ*| ≫ 1 but *N_W_*ϵ*^2^* ≪ 1, and whatever the sign of *ϵ* in the regime where *ϵ* ≪ 1 and *N_W_* |*ϵ*| ≪ 1, but there is suppression for *ϵ* > 0 satisfying *ϵ* ≪ 1 and *N_W_* |*ϵ*| ≫ 1 but *N_W_*ϵ*^2^* ≪ 1.

Interestingly, in the regime of very small mutational effects where *ϵ* ≪ 1 and *N_W_* |*ϵ*| ≪ 1, we showed that the strongest amplification is obtained in the limit *α* ≫ 1, but now we find that in this case, amplification is only transient. Our expansions show that universal amplification can exist only if 1 < *α* ≤ *D* − 1. In this case, for *ϵ* ≪ 1 and *N_W_* |*ϵ*| ≪ 1, the strongest amplification is expected for *α* = *D* − 1 (because *F* increases with *α*, see Eq. S40), and we then have *F* (*D* − 1) = (*D* − 2)^3^/{*D* [*D* (*D* − 2) + 2]} so that Eq. S39 then yields

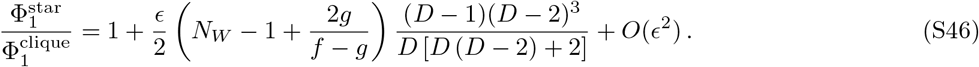

If *N_W_* ≫ 1 and *D* ≫ 1, this gives a prefactor of the first order term in *ϵ* of order *N_W_ D*/2, which can yield a large amplification, but recall that this is restricted to *N_W_* |*ϵ*| ≪ 1.

#### 4. Expansion for extremely asymmetric migrations

So far we have considered expansions in selection strengths, and then in some regimes, analyzed extremely asymmetric migrations. However, the order of limits matters and our previous discussions are limited to specific regimes in terms of selection strength. If *α* → 0, Eq. S37 yields

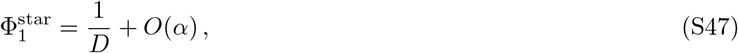

which demonstrates that for small *m* the star is a very strong suppressor of selection, with all mutations becoming effectively neutral (once they have fixed in a deme). Note that this is consistent with our result for very small mutational effects, see Eq. S41 and the discussion just below.

If *α* → ∞, and in particular assuming *α* ≫ *D*, Eq. S37 yields

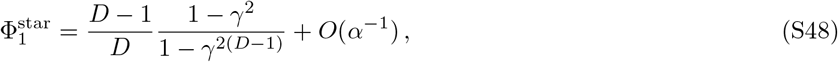

and for *γ* = *N_W_ ρ_W_*/(*N_M_ ρ_M_*) → 0, which occurs when *f_M_* ≫ *f_W_*, we have 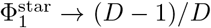, which confirms that amplification can only be transient in this case, since for the clique, we have 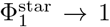 in this limit. The simple expression 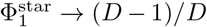 is due to the fact that in the *α* → ∞, mutants in the center cannot fix even if they are very fit, while those in the leaves fix easily. If in addition *D* ≫ 1 (and thus *α* ≫ *D* ≫ 1), then

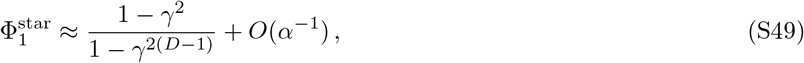

which is formally reminiscent of the fixation probability for the star in the model of Ref. [4] with Birth-death dynamics in the limit *D* → ∞ (see also Ref. [28]), where *γ* replaces the ratio of fitnesses (as is the case in Eq. 2 for the clique, which is formally reminiscent of the fixation probability for the well-mixed population). Thus, if *α* → ∞ such that *α* ≫ *D* ≫ 1, the star can become a universal amplifier of selection with respect to the clique, and has a fixation probability identical to that of the clique but with *γ* replaced by *γ*^2^, which demonstrates amplification, in the same way as in the model of Ref. [4]. However, this is restricted to the particular regime *α* → ∞ such that *α* ≫ *D* ≫ 1.

In the Birth-death model [4], the star satisfies *m_O_* = 1/(*D* − 1) and *m_I_* = 1, so that *α* = *D* − 1 (see section III for a more general mapping between our model and that of Ref. [4]). Let us thus consider the specific case where *α* = *D* − 1. If *α* → ∞ (which implies *D* → ∞), Eq. S37 yields

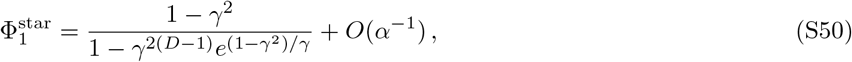

which has the exact same form as the rigorous asymptotic expression [29] for *D* → ∞ of the fixation probability in a star in the Birth-death model of Ref. [4]. This is consistent with the fact that Eqs. S34, S36 and S37 are exactly equivalent to the formula given in Ref. [28] with *γ* → *f_W_*/*f_M_* and *α* → *D* − 1 (see above). Note that the rigorous asymptotic expression from Ref. [29] is slightly different from the better known expression that has the same form as Eq. S49, which holds for *α* ≫ *D* ≫ 1 in our model.

#### 5. Additional results for the star

In Fig. S2, we show results for the fixation probability in the star graph that complement those shown in Fig. 2.

**FIG. S2.**
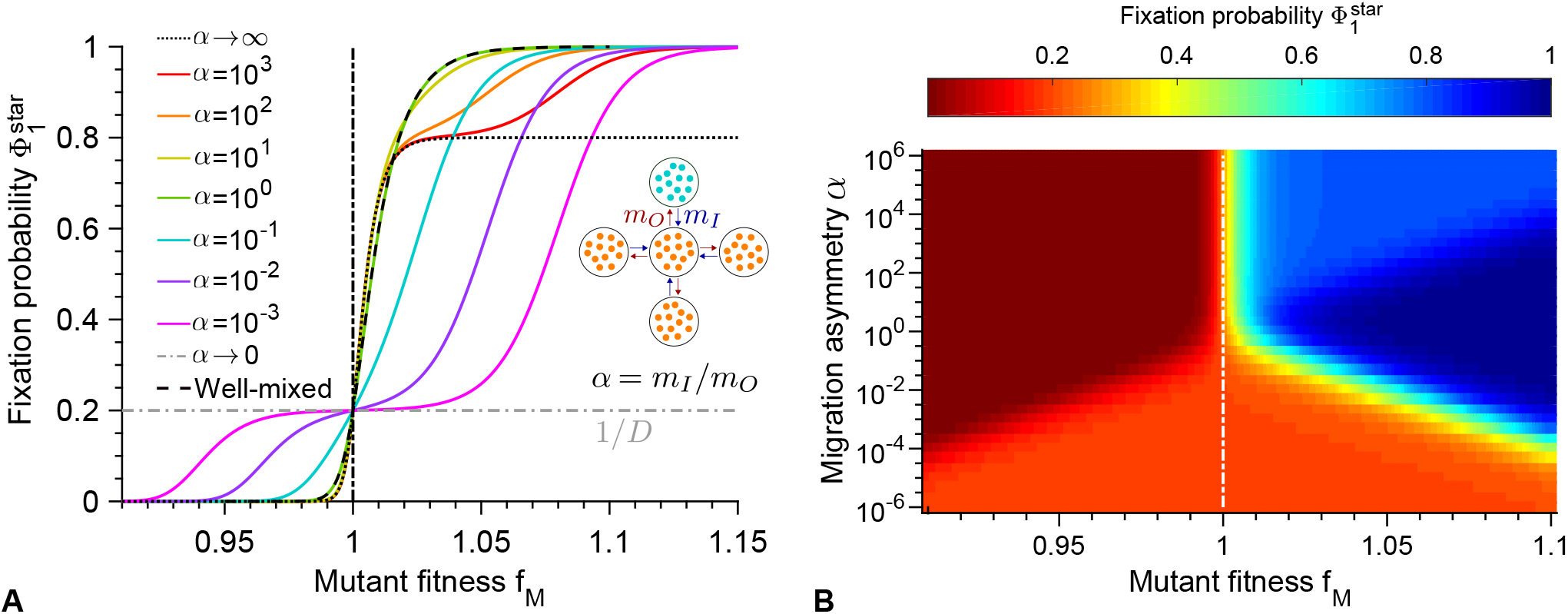
Fixation probability for the star. **A:** Fixation probability 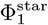 of mutants in a star graph versus mutant fitness *f_M_*, starting with one fully mutant deme chosen uniformly at random, with different migration rate asymmetries *α* = *m_I_/m_O_*, complementing those shown in Fig. 2A. Data for the well-mixed population is shown as reference, with same total population size and initial number of mutants. Curves represent analytical predictions in Eq. S37. **B:** Heatmap of the same fixation probability 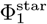 shown versus mutant fitness *f_M_* and migration rate asymmetry *α* = *m_I_*/*m_O_*. Parameter values in both panels: *D* = 5, *K* = 100, *f_W_* = 1, *g_W_* = *g_M_* = 0.1. Vertical dash-dotted lines represent the neutral case *f_W_* = *f_M_*.

In this work, we usually start from a mutant deme chosen uniformly at random, which is realistic for spontaneous mutations. However, the initial position of the mutant [4], and the degree of the node where it starts [11], can strongly impact its fate. Thus, in Figs. S3 and S4, we show results when the initial mutant deme is either the center or a leaf. These results illustrate the strong impact of mutant initial position.

**FIG. S3.**
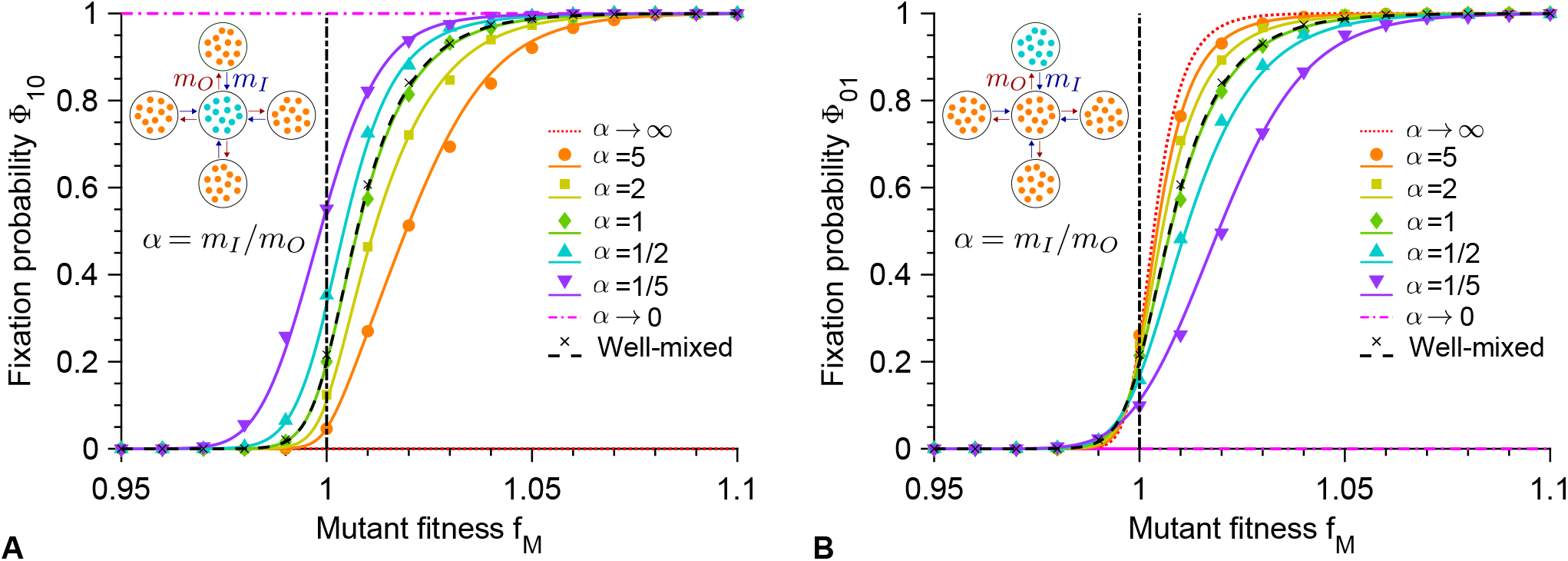
Fixation probability for the star, starting from a mutant center or leaf. **A:** Fixation probability Φ_10_ of mutants in a star graph versus mutant fitness *f_M_*, starting with a fully mutant center, with different migration rate asymmetries *α* = *m_I_*/*m_O_*. Markers are computed over 10^3^ stochastic simulation realizations. Curves represent analytical predictions in Eq. S34. **B:** Fixation probability Φ_01_ of mutants in a star graph versus mutant fitness *f_M_*, starting with a fully mutant leaf, with different migration rate asymmetries *α* = *m_I_*/*m_O_*. Markers are computed over 10^3^ stochastic simulation realizations. Curves represent analytical predictions in Eq. S36. In both panels, vertical dash-dotted lines represent the neutral case *f_W_* = *f_M_*. Data for the well-mixed population is shown as reference, with same total population size and initial number of mutants. Parameter values in both panels: *D* = 5, *K* = 100, *f_W_* = 1, *g_W_* = *g_M_* = 0.1. From top to bottom, (*m_I_, m_O_*) 10^6^ = (5, 1); (2, 1); (1, 1); (1, 2); (1, 5) in simulations.

**FIG. S4.**
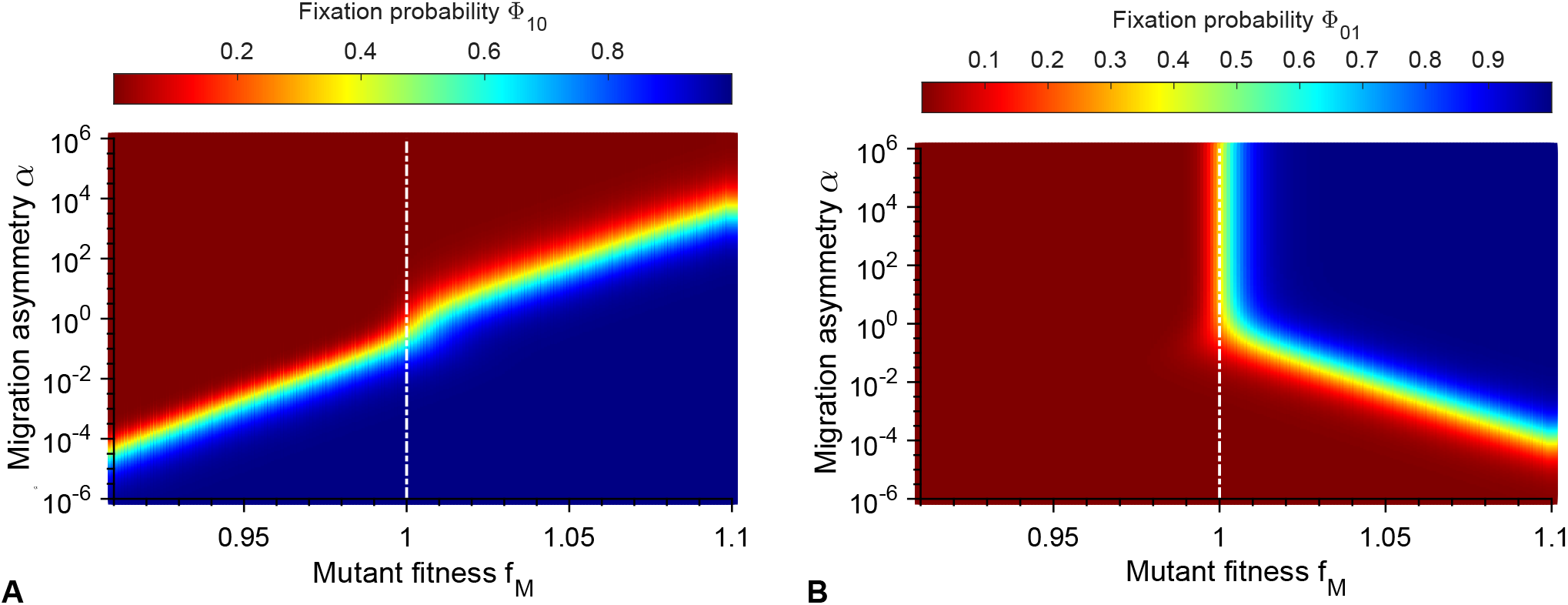
Heatmaps of the fixation probability for the star, starting from a mutant center or leaf. **A:** Heatmap of the fixation probability Φ_10_ of mutants in a star graph starting with a fully mutant center, shown versus mutant fitness *f_M_* and migration rate asymmetry *α* = *m_I_*/*m_O_*. **B:** Similar heatmap but for the fixation probability Φ_01_ of mutants in a star graph starting with a fully mutant leaf. In both panels, vertical dash-dotted lines represent the neutral case *f_W_* = *f_M_*. Parameter values in both panels: *D* = 5, *K* = 100, *f_W_* = 1, *g_W_* = *g_M_* = 0.1.

## III. COMPARISON WITH THE MODEL OF REF. [4]

In Ref. [4], a model where each of the *N* nodes of a graph is occupied by a single individual was introduced. Replacement probabilities *w_ij_* from node *i* to node *j* are defined along each edge *ij* of the graph. At each elementary step, an individual (say the one on node *i*) is selected for division, with probability proportional to fitness *f_i_*, and its offspring replaces the individual on node *j* with probability *w_ij_*. This dynamics, which became known as the Birth-death dynamics [12–14] or biased invasion process [11, 16], thus allows to always maintain exactly one individual on each node. An important constraint stemming from the definition of the model is

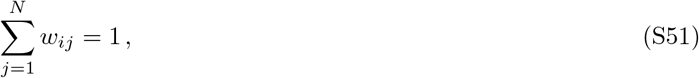

because the offspring of individual *i* has to end up somewhere. In other words, the matrix of replacement probabilities is right-stochastic. Note that self-loops where the offspring stays on the same node (corresponding to *w_ii_* > 0) were not considered in the initial description of the model but can be added (see e.g. [23]). The probability *P_i_*_→*j*_ that, at a given elementary step, the offspring from node *i* replaces the individual in node *j* is given by

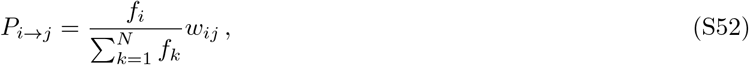

i.e. the probability that the individual on node *i* is selected for division, multiplied by the probability that its offspring replaces the individual in node *j*. Note that, since exactly one replacement occurs per elementary step, Σ_*i,j*_ *P_i→j_* = 1, and that Eq. S52 satisfies this normalization constraint because Eq. S51 holds. Using Eq. S51, we can rewrite Eq. S52 as

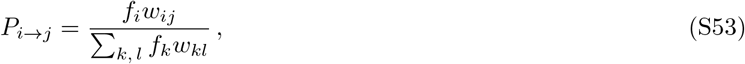

which will be convenient for our comparison.

In our coarse-grained model, upon each migration event, the individual that migrated from deme *i* to deme *j* (with migration rate *m_ij_* per individual) may fix with probability *ρ_i_*. In particular, the probability 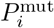 that a specific deme *i* becomes mutant upon one given migration event while it was wild-type before reads

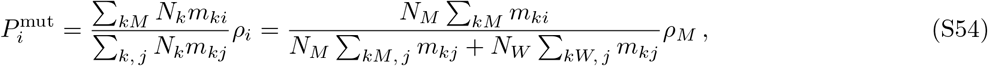

where Σ_*kM*_ denotes a sum over mutant (*M*) demes indexed by *k*. In the last term we discriminated over mutant and wild-type demes and employed the fact that all demes have the same carrying capacity *K*, resulting in steady-state sizes *N_M_* for mutant demes and *N_W_* for wild-type demes, and denoted by *ρ_M_* the fixation probability of a mutant in a wild-type deme, following our usual convention. Here, we have considered a probability upon a migration event, but migration events change the makeup of the population only if fixation ensues. To compare to the model of Ref. [4], let us instead focus only on the migration events that result into fixation. The probability 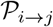 that, upon such a successful migration event, an individual coming from deme *i* fixes in deme *j* reads

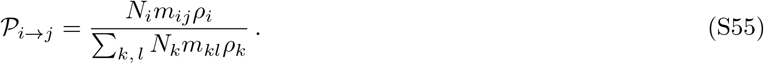

Note that it satisfies 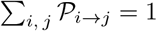, as a fixation occurs at each successful migration event.

Eqs. S53 and S55 have the same form, with 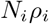 in our model playing the part of *f_i_* in the model of Ref. [4]. An important difference is that in our model, the *m_ij_* (which are migration rates, not migration probabilities) do not need to satisfy the constraint in Eq. S51 and are independent. Our model is thus less constrained than that of Ref. [4]. Note that an alternative dynamics removing this constraint was discussed in Ref. [4], but then very rarely considered in the literature [14]. Note also that the fixation probability *ρ_i_* involves the fitness of *i* and that of the type that is replaced (say *j*), but because we always work with just two types, this dependence can be ignored without losing generality.

For the clique and for the cycle, we have found that the fixation probability, given by Eq. S13, has the same form as that for the well-mixed population, but with 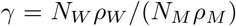 playing the part of *f_W_*/*f_M_*. This is perfectly consistent with the mapping described here, with 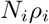 in our model playing the part of *f_i_*. Note that the constraint on migration rates does not come into play here since Eq. S13 is independent of migration rates.

For the star, we already noted that the fixation probabilities given in Eqs. S34, S36 and S37 are exactly equivalent to the formula given in Ref. [28] if the following substitutions are made: 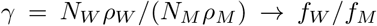 and *α* = *m_I_*/*m_O_* → *D* − 1 (bearing in mind that in the notations of Ref. [28], *D* − 1 is called *n* and *f_W_*/*f_M_* is called 1/*r*). Again, this is perfectly consistent with the mapping described here, with 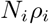 in our model playing the part of *f_i_*. In addition, we have to impose a specific value of *α*, namely *α* = *D* − 1, in order to get back the result of Ref. [28]. This is because, for a star graph with no self-loops, only two different migration rates can exist, *m_O_* from center to leaf and *m_I_* from leaf to center, given the symmetries of this graph. Imposing that Σ_*j*_ *m_ij_* is independent on *i*, i.e. that all nodes have the same total emigration rate, which is a weaker form of the constraint in Eq. S51 (because no normalization is required on migration rates), then yields *α* = *D* − 1. So the extra constraint in the mapping between the two models for the star stems from the requirement that Eq. S51 be satisfied in the model of Ref. [4]. This also means that for *α* = *D* − 1, our results are formally the same as in Ref. [28] and in the model of Ref. [4] and that we then find the exact same amplification properties for the star. But this exact correspondence is restricted to a very particular value of *α*.

## IV. GENERALIZED CIRCULATION THEOREM

Here, we extend the circulation theorem from Ref. [4] to our model. Consider a metapopulation on a graph *G* with a set of nodes ***V*** where all *D* demes have the same carrying capacity. A graph with migration rates per individual *m_ij_* from node *i* to node *j* is a circulation if and only if for all *i*,

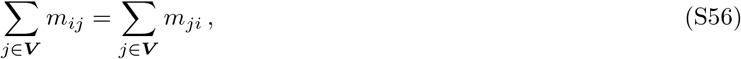

which means that the total rate of migrations leaving *i* is equal to the total rate of migrations arriving in *i*. We will show that the fixation probability starting from *P* fully mutant demes is the same as for the clique, i.e. is given by Eq. S13, if and only if the graph *G* is a circulation. Let us denote by ***P*** the ensemble of fully mutant demes and by *P* its cardinal.

Following the proof of the circulation theorem given in Ref. [4], we will demonstrate that the following are equivalent:

1. G is a circulation.
2. *P* performs a random walk with forward bias *γ*^−1^, with 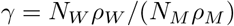, and absorbing states {0, *D*}.
3. The fixation probability starting from *P* fully mutant demes is the same as for the clique, i.e. is given by Eq. S13.
4. The probability that, starting from any *P* fully mutant demes, a mutant such that 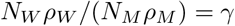 eventually fixes in *P′* mutant demes is given by

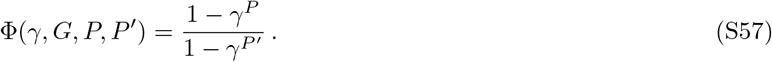

First we show that (1) ⇒ (2), in a similar way as in Ref. [4]. For this, let *δ*_+_(***P***) (resp. *δ*_−_(***P***)) be the probability that the number of mutant demes increases by one (resp. decreases by one). We have

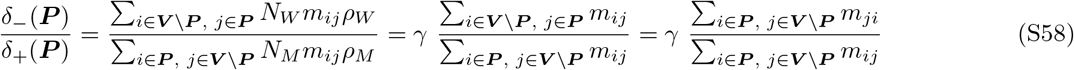

Since *G* is a circulation, Eq. S56 holds, and summing it over all *i* ∈ ***P*** yields

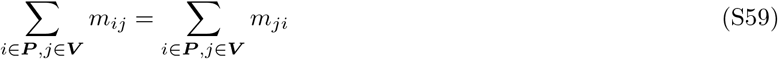

which can be rewritten as

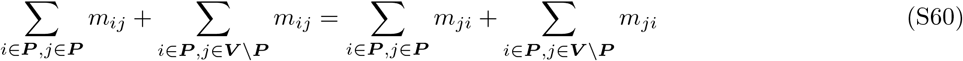

and thus

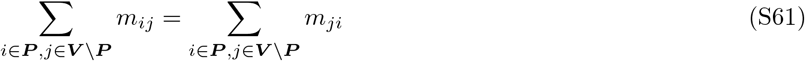

so that Eq. S58 becomes

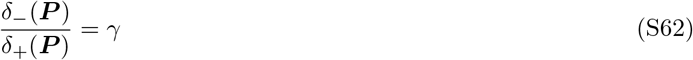

and thus *P* performs a random walk with forward bias *γ*^−1^.

(2) ⇒ (3) can be proved as for the clique (see section II A and Ref. [25]).

(3) ⇒ (4) can be proved using conditional probabilities exactly as in Ref. [4].

(4) ⇒ (1) can also be proved similarly as in Ref. [4]. Specifically, using Eq. S57 for *P* = 1 and *P′* = 2 gives

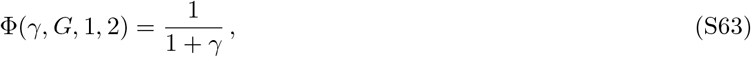

but denoting by *v* the initially mutant deme, we can also write the probability that 2 demes become mutant after any number *k* of migration events as

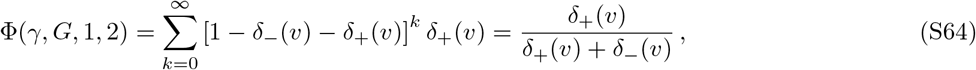

and comparing Eqs. S63 and S64 shows that for any initially mutant deme *v*,

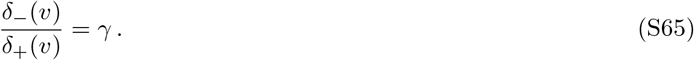

But Eq. S58 yields

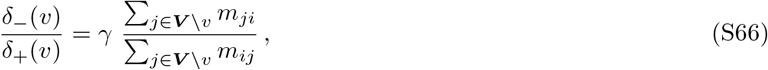

and therefore, for all *v*,

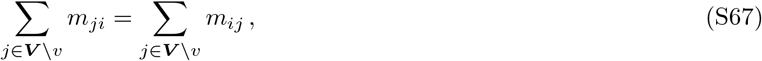

which entails

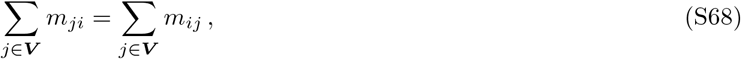

and thus *G* is a circulation (see Eq. S56).

## V. COMPARISON WITH THE MODEL OF REF. [16]

In Ref. [16], a model generalizing that of Ref. [4] to the case where each node of the graph is occupied by a deme with a fixed number of individuals was introduced. In the models of Refs. [4] and [16], each elementary event is composed of a death event in one deme and a birth event in another one, thus allowing to maintain constant the population of each deme. Furthermore, the order employed to choose the individual that dies and the one that divides matters for final results, yielding Birth-death and death-Birth models, as in the model introduced in Ref. [4] (see Refs. [12–14]). Conversely, in our model, migration, death and birth events are all independent. This is made possible by allowing the population size of each deme to vary. Here, we present the model of Ref. [16] and compare it to our model.

Let us consider wild-type fitness as reference and set it to 1, and let us denote mutant fitness by 1 + *s*. Let us denote the total number of individuals in deme *i* by *N_i_*, and the number of mutant individuals in deme *i* by *n_i_*. Migration probabilities *w_ij_* from deme *i* to deme *j* are defined along each edge *ij* of the graph. We will denote by 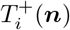 the transition probability from *n_i_* to *n_i_* + 1 and by 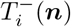 the transition probability from *n_i_* to *n_i_* − 1, which both depend on the complete state of the system ***n*** = (*n*_1_, *n*_2_,…, *n_M_*).

### 1. Birth-death dynamics

In Birth-death dynamics (also known as “biased invasion process” [11, 16]), the *w_ki_* satisfy the normalization constraint

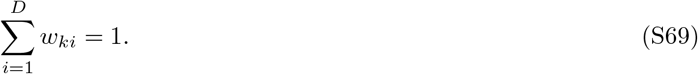

In this dynamics, an individual is chosen for reproduction among all the individuals of the population according to its fitness. Assuming that it belongs to island *k*, its offspring migrates to island *i* with probability *w_ki_*, where it replaces an individual chosen uniformly at random among the *N_i_* individuals there. The transition probability 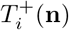 is given by

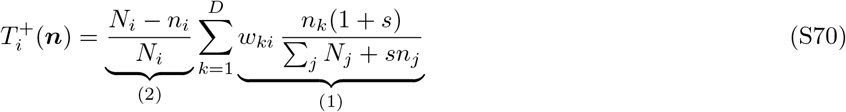

where (1) is the probability for a mutant to reproduce on island *k* and to migrate to *i*, which is then summed over all the islands *k*, and (2) is the probability that, given that a death event occurs on island *i* (because an individual in *i* is being replaced), a wildtype individual dies. Analogously:

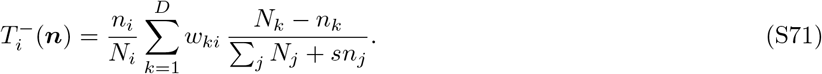

### 2. Death-birth dynamics

In death-Birth dynamics (also known as “biased voter model” [11, 16]), we assume that Σ_*k*_ *w_ki_* = 1. In this dynamics, an individual is chosen uniformly at random in the entire population to die. Assuming that death occurred in island *i*, one may consider that a migration event then occurs from island *k* to *i* with probability *w_ki_*. But one may also assume that migration occurs from *k* to *i* with a probability proportional to the product of *w_ki_* and the total fitness *N_k_* + *sn_k_* of island *k*. The first choice considers fitness to be relevant only within each island, while the second one takes into account fitness across the islands. We will consider the second one because it allows to recover the usual death-Birth model [12–14] when *N_i_* = 1 ∀*i* ∈ {1, …, *D*}. Finally, the reproducing individual on island *i* is chosen according to its fitness within the island. The transition probability 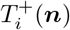 reads

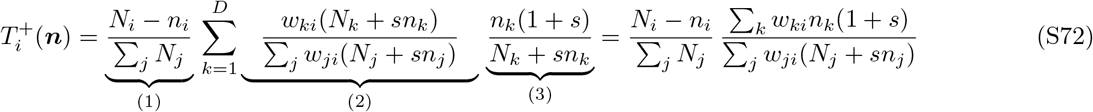

where (1) is the probability for a wildtype individual to die on island *i*, while (2) is the probability that, given that a death event occurs on island *i*, a migration event occurs from island *k* to *i*, which takes into account the total fitness of island *k*. Finally, (3) is the probability that, given that a reproduction event happens in island *k*, it is a mutant who reproduces. Similarly,

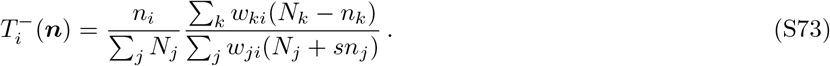

### 3. Clique

Consider a clique made of *D* demes of size *N* (all of identical and composition-independent size), such that *w_ij_* = *w* for all *i* ≠ *j* and *w_ii_* = *w′* for all *i*. If migrations between different demes are rare enough, one can coarse-grain the process and consider that each deme is either fully mutant or fully wild-type, and the state of the clique can then be fully described by the number *ξ* of mutant demes, which changes when migrations followed by fixation occur. In the Birth-death model, starting from Eq. S71 and summing over mutant demes, we obtain

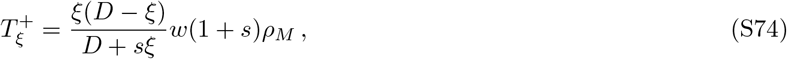

and similarly

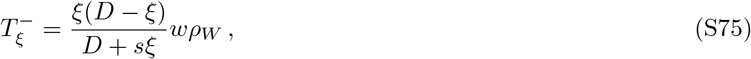

which entails that

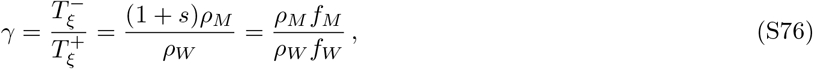

and thus the probability that the mutant fixes in the whole population starting from *i* mutant demes is given by Eq. S13 but with *γ* expressed in Eq. S76. The same result is obtained in the death-Birth case.

### 4. Star

We consider the star graph with self-loops (i.e. allowing replacement within a given deme, corresponding to migration from this deme to itself, *w_ii_* > 0). Indeed, the rare migration regime that we study in our model means in the framework of the model of Ref. [16] that replacement is much more frequent within a deme than across two demes, thus requiring very strong self-loops, i.e. large *w_ii_* values. While the star with self-loops was introduced in Ref. [23] with one individual per node of the graph, in the spirit of Ref. [4], here we treat it in the model of Ref. [16], where each node contains a deme with fixed size *N*. Following Ref. [23], we introduce two parameters *x* and *y* (0 < *x*, *y* ≤ 1) such that 1 − *y* is the weight of the self-loop of the center and 1 − *x* is the weight of the self-loops on each leaf. The other weights are chosen in order to respect the symmetry of the star and for *W* to be right stochastic in the Birth-death case and left stochastic in the death-Birth case. Hence, in the Birth-death model, the matrix of migration probabilities reads:

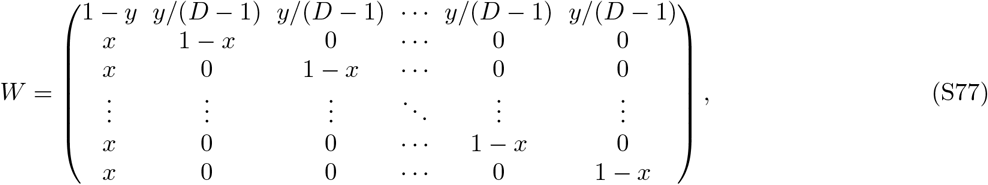

where nodes are numbered so that the first one is the center of the star and others are leaves. The case *x* = *y* = 1 corresponds to the star without self-loops introduced in Ref. [4].

In order to compare our model to the model of Ref. [16] in the case of the star, we choose their respective parameters so that in each deme, both models have the same value for the ratio of the migration rate *T_mig_* leaving the deme to the reproduction rate *T_rep_* in the same deme. In our model, the reproduction rate per individual is given by *T_rep_* = *f_W_* (1 − *N_W_*/*K*) for each deme, whatever its type (leaf or center) – in the wild-type case. Still in our model, the migration rate leaving the center (to any leaf) is *m_O_*(*D* − 1) per individual, and that leaving a leaf (to the center, which is the only possibility) is *m_I_* per individual. In the framework of the Birth-death model of Ref. [16], the total reproduction probability per individual in a deme (irrespective of where the offspring from this deme migrates) is equal to Σ_*j*_ *w_ij_* = 1, for both the center and for a leaf, while the total migration probability per individual leaving the center is Σ_*j*≠1_ *w*_1*j*_ = *y* and the one leaving a leaf is Σ_*j*≠*i*_ *w_ij_* = *x* with *i* > 1 (see Eq. S77). Thus, to match our model with the Birth-death model of Ref. [16], we have the following two constraints:

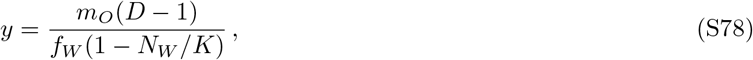

and

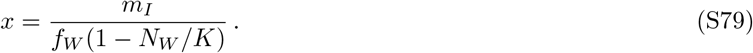

Fig. S5A shows that once this matching is done, a good agreement is obtained between simulation results for the two models, which yield similar mutant fixation probabilities across various migration asymmetries *α*. Figs. S5B and C show small relative and absolute differences, respectively, between the two models. Note that the relative error is high when the probability of fixation of a mutant deme is close to zero, which is the case for deleterious mutations, but then the absolute error is small, which confirms that these models are consistent.

**FIG. S5.**
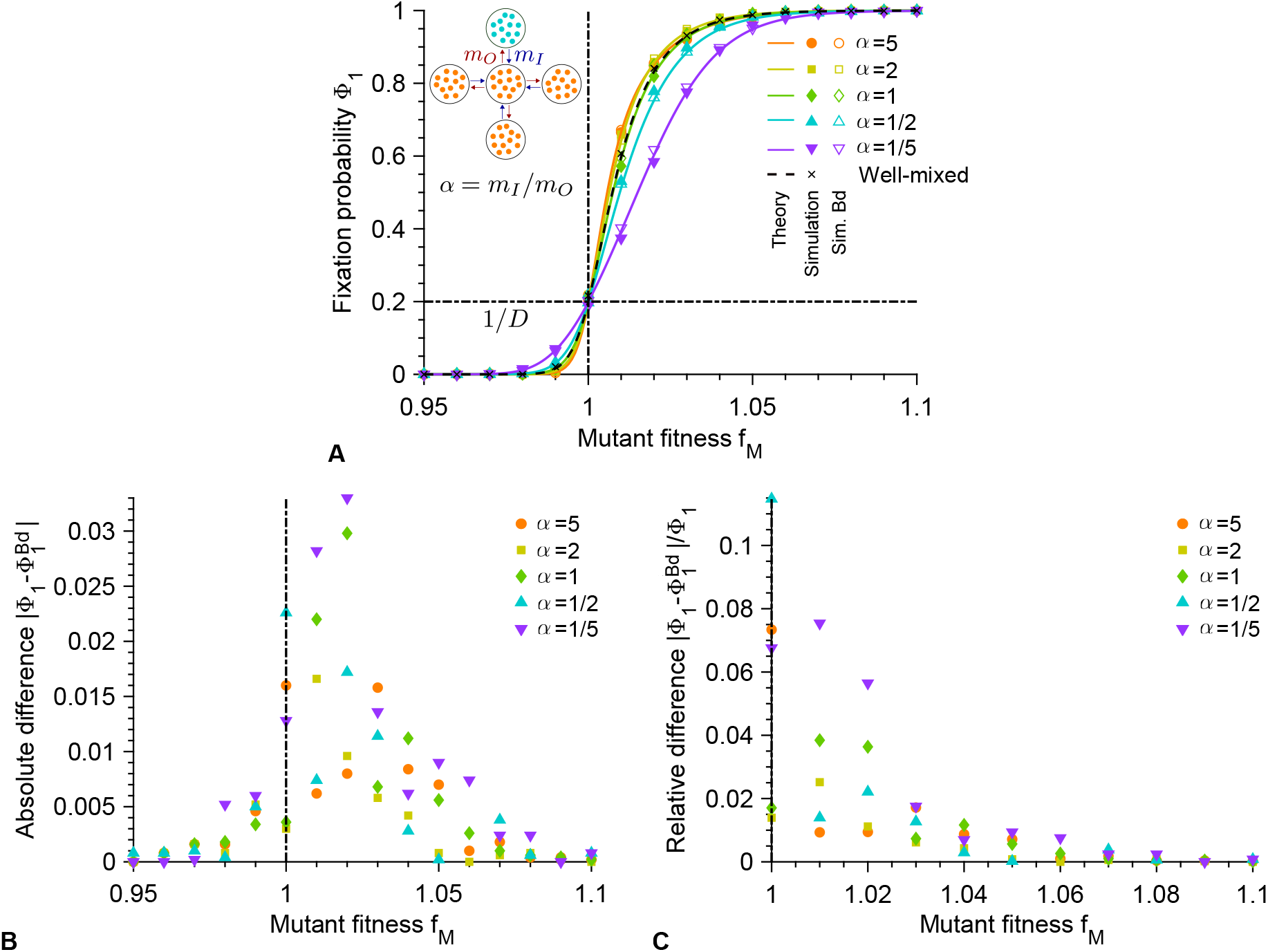
Comparison between the Birth-death model inspired by Refs. [16] and [23] and our model. **A:** Fixation probability Φ_1_ of mutants in a star graph versus mutant fitness *f_M_*, starting with one fully mutant deme chosen uniformly at random, with different migration rate asymmetries *α* = *m_I_*/*m_O_* in our model and in the matching Birth-death (Bd) model, which satisfies Eqs. S78 and S79. Markers are obtained from 2 × 10^3^ stochastic simulation realizations in our model, and in the Bd model of Ref. [16]. Curves represent analytical predictions for our model in Eqs. S34, S36 and S37. **B:** Absolute differences between simulation results obtained with the two models (see panel **A**), as a function of the mutant fitness *f_M_*. **C:** Relative differences between simulation results obtained with the two models (see panel **A**), as a function of the mutant fitness *f_M_*. Parameter values for our model: *D* = 5, *K* = 100, *f_W_* = 1, *g_W_* = *g_M_* = 0.1, and from top to bottom in the legend of panel **A**, (*m_I_*, *m_O_*) × 10^6^ = (5, 1); (2, 1); (1, 1); (1, 2); (1, 5) in simulations, as in Fig. 2. Parameter values for the matching Birth-death model: *D* = 5, *N* = *N_W_* = 90, *f_W_* = 1, and values of *x* and *y* satisfying Eqs. S78 and S79 for each pair of values of *m_I_* and *m_O_* from our model. Vertical dash-dotted lines indicate the neutral case *f_W_* = *f_M_*.

The Birth-death model of Ref. [16] has the same total reproduction rate in each deme. Once the matching in Eqs. S78 and S79 is done, it also features the same migration-to-reproduction ratio as in our model. Note however that the death rate is not uniform across demes in this model: in the center it is Σ_*i*_ *w*_*i1*_ = 1 − *y* + (*D* − 1)*x*, and in a leaf it is Σ_*i*_ *w*_*ij*_ = 1 − *x* + *y*/(*D* − 1) with *j* > 1. This stands in contrast with our model, and to resolve this discrepancy, we would need to impose that *y* = (*D* − 1)*x*. In that case, the matrix of migration probabilities in Eq. S77 becomes doubly stochastic and the star becomes a circulation, and thus it has the same fixation probability as the clique in the model of Ref. [16] (see above). Consistently, Eqs. S78 and S79 then entail *α* = 1. This shows that the matching between models is not perfect for other values of *α*, because the model of Ref. [16] is more constrained than our model, as it imposes constant deme size.

In the death-Birth model, the matrix of migration probabilities is the transpose of that given in Eq. S77. Hence, the total reproduction rates for a leaf and the center are Σ_*i*_ *w*_ij_ = *y*/(*D*−1)+1−*x* with *j* > 1 and Σ_*i*_ *w*_*i1*_ = 1−*y*+(*D*−1)*x*, respectively, while the total migration rates per individual from a leaf and from the center are *y/*(*D* − 1) and (*D* − 1)*x*, respectively. Thus, to match our model with the death-Birth model, we have the following two constraints:

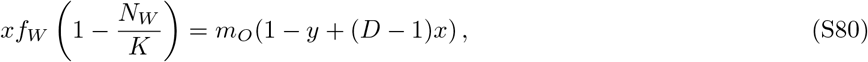

and

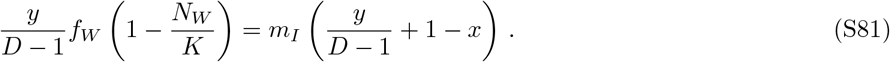

In this case too, Fig. S6 shows that once this matching is done, a good agreement is obtained between simulation results for the two models.

**FIG. S6.**
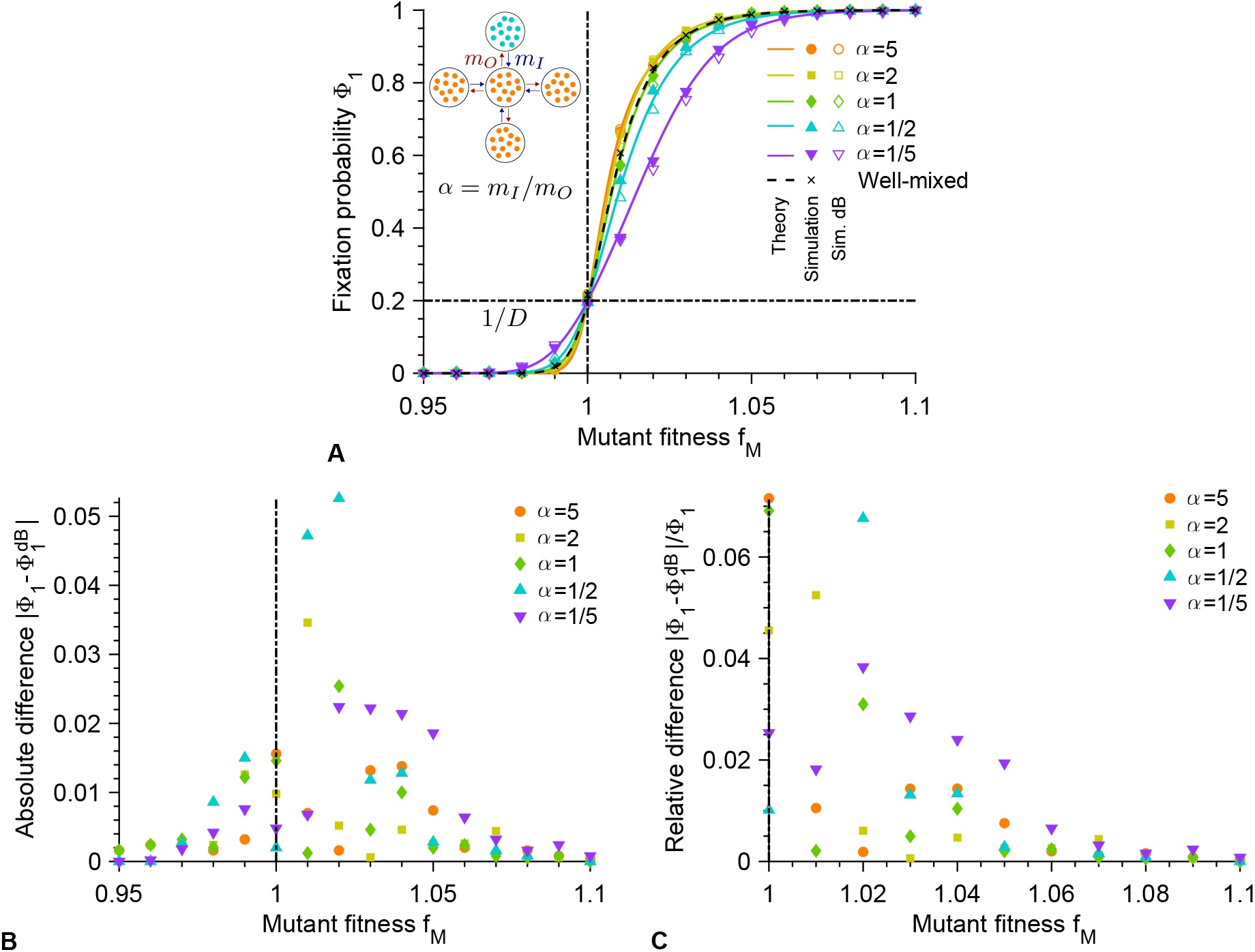
Comparison between the death-Birth model inspired by Refs. [16] and [23] and our model. **A:** Fixation probability Φ_1_ of mutants in a star graph versus mutant fitness *f_M_*, starting with one fully mutant deme chosen uniformly at random, with different migration rate asymmetries *α* = *m_I_*/*m_O_* in our model and in the matching death-Birth (dB) model, which satisfies Eqs. S80 and S81. Markers are obtained from 2 × 10^3^ stochastic simulation realizations in our model, and in the dB model of Ref. [16]. Curves represent analytical predictions for our model in Eqs. S34, S36 and S37. **B:** Absolute differences between simulation results obtained with the two models (see panel **A**), as a function of the mutant fitness *f_M_*. **C:** Relative differences between simulation results obtained with the two models (see panel **A**), as a function of the mutant fitness *f_M_*. Parameter values for our model: *D* = 5, *K* = 100, *f_W_* = 1, *g_W_* = *g_M_* = 0.1, and from top to bottom in the legend of panel **A**, (*m_I_*, *m_O_*) × 10^6^ = (5, 1); (2, 1); (1, 1); (1, 2); (1, 5) in simulations, as in Fig. 2. Parameter values for the matching death-Birth model: *D* = 5, *N* = *N_W_* = 90, *f_W_* = 1, and values of *x* and *y* satisfying Eqs. S80 and S81 for each pair of values of *mI* and *mO* from our model. Vertical dash-dotted lines indicate the neutral case *f_W_* = *f_M_*.

The death-Birth model of Ref. [16] has the same total death rate in each deme. Once the matching in Eqs. S78 and S79 is done, it also features the same migration-to-reproduction ratio as in our model. Note however that the birth rate is not uniform across demes in this model even in the absence of fitness differences (see above). Here too, to resolve this discrepancy with our model, we would need to impose that *y* = (*D* − 1)*x*, with the same consequences as in the Birth-death model – note that Birth-death and death-Birth models then yield the same result. Again, this shows that the matching between models is not perfect for other values of *α*, because the model of Ref. [16] is more constrained than our model, as it imposes constant deme size.

## VI. EXTENSION TO DIFFERENT DEME SIZES: THE DOUBLET

### A. Main results

Our model allows us to consider structures involving demes with different sizes. In this case, we consider an initial mutant placed randomly with a probability proportional to deme size, which is realistic for mutations occurring upon division or with a constant rate per individual (note that this corresponds to both uniform and temperature initial conditions in the language of models with a single individual per node [23], which coincide in our model).

As a simple example, consider a doublet comprising a small deme with carrying capacity *K*_*S*_ and a larger deme with carrying capacity *K*_*L*_ > *K*_*S*_ (see Fig. S1D). Individuals can migrate from the large (resp. small) deme to the small (resp. large) deme with a rate per individual *m*_*S*_ (resp. *m*_*L*_). For structured populations involving demes with identical sizes, we considered the fixation probability Φ_1_ starting from one fully mutant deme, which yields that of one mutant individual when multiplied by *ρ*_*M*_. Here, we consider the fixation probability of one single mutant in the structure divided by that in the small deme. If we define *D* such that *K*_*L*_ = (*D* − 1)*K*_*S*_ and if we choose the notation *K*_*S*_ = *K*, then this quantity 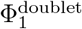 is analogous to our usual Φ_1_ (if *D* is an integer), thus facilitating comparisons. 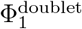 is expressed analytically below.

Fig. S7 shows 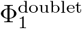 for different migration asymmetries *α* = *m*_*S*_/*m*_*L*_, with excellent agreement between our analytical predictions and our simulation results (see also Fig. S8 for additional *α* values and a heatmap). Furthermore, Fig. S7 shows that the fixation probability is very close to the well-mixed case when *α* = 1/(*D* − 1). This corresponds to *m*_*S*_*K*_*L*_ = *m*_*L*_*K*_*S*_, i.e. to equal migration flows from small to large deme and reciprocally. We also observe that the doublet behaves as a suppressor of selection for *α* > 1/(*D* − 1), and has weak amplifying properties for *α* < 1/(*D* − 1), which do not survive in the limit *α* → 0. In the Appendix, Section VI, we show that in the regime of moderate mutational effects, the doublet is an amplifier of selection with respect to the clique for 1/(*D* − 1)^2^ < *α* < 1/(*D* − 1), and a suppressor of selection for *α* > 1/(*D* − 1). This generalizes the result of Ref. [4] that small upstream populations with large downstream populations, corresponding here to *α* → 0, yield suppressors. Furthermore, this confirms the importance of migration asymmetry in the impact a population structure has on selection.

**FIG. S7.**
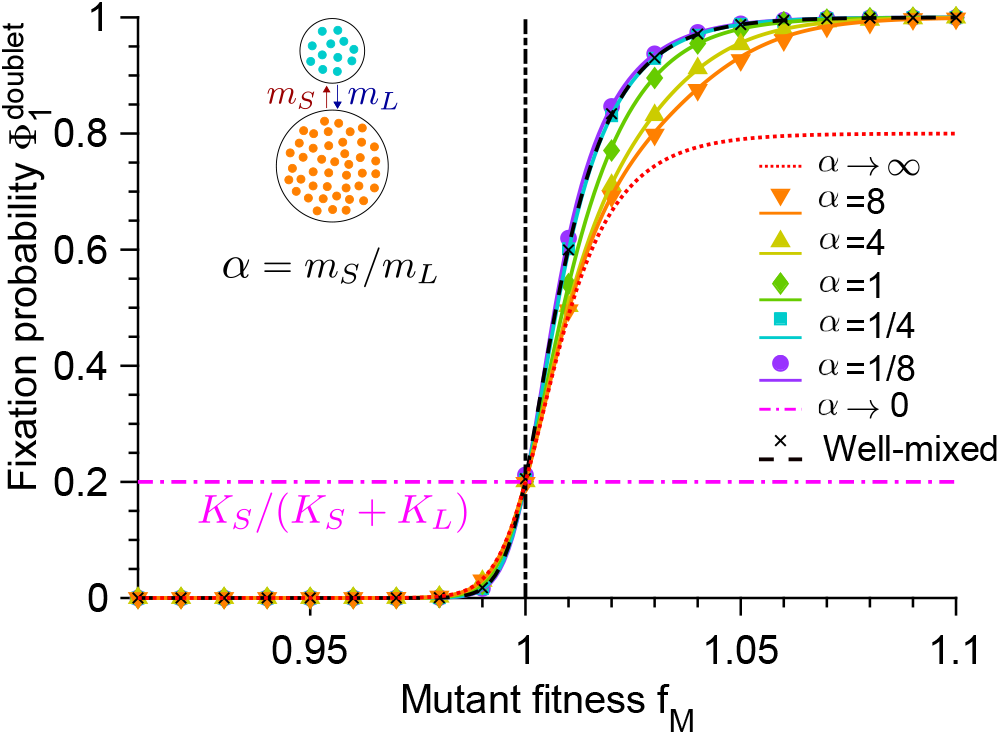
Fixation probability for the doublet. Fixation probability 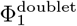 of mutants in a doublet versus mutant fitness *f*_*M*_, starting with one fully mutant deme chosen proportionally to deme size, with different migration asymmetries *α* = *m*_*S*_/*m*_*L*_. Data for the well-mixed population is shown as reference, with same total population size and initial number of mutants. Markers are computed over 2 × 10^3^ stochastic simulation realizations. Curves represent analytical predictions in Eqs. S84, S85 and S87. Vertical dash-dotted lines indicate the neutral case *f*_*W*_ = *f*_*M*_. Parameter values: *K*_*S*_ = 100, *K*_*L*_ = 400 (hence *K*_*L*_ = (*D* − 1)*K*_*S*_ with *K*_*S*_ = *K* = 100 and *D* = 5), *f*_*W*_ = 1, *g*_*W*_ = *g*_*M*_ = 0.1. From top to bottom,(*m*_*S*_, *m*_*L*_)× 10^6^ = (8, 1); (4, 1); (1, 1); (1, 4); (1, 8) in simulations.

**FIG. S8.**
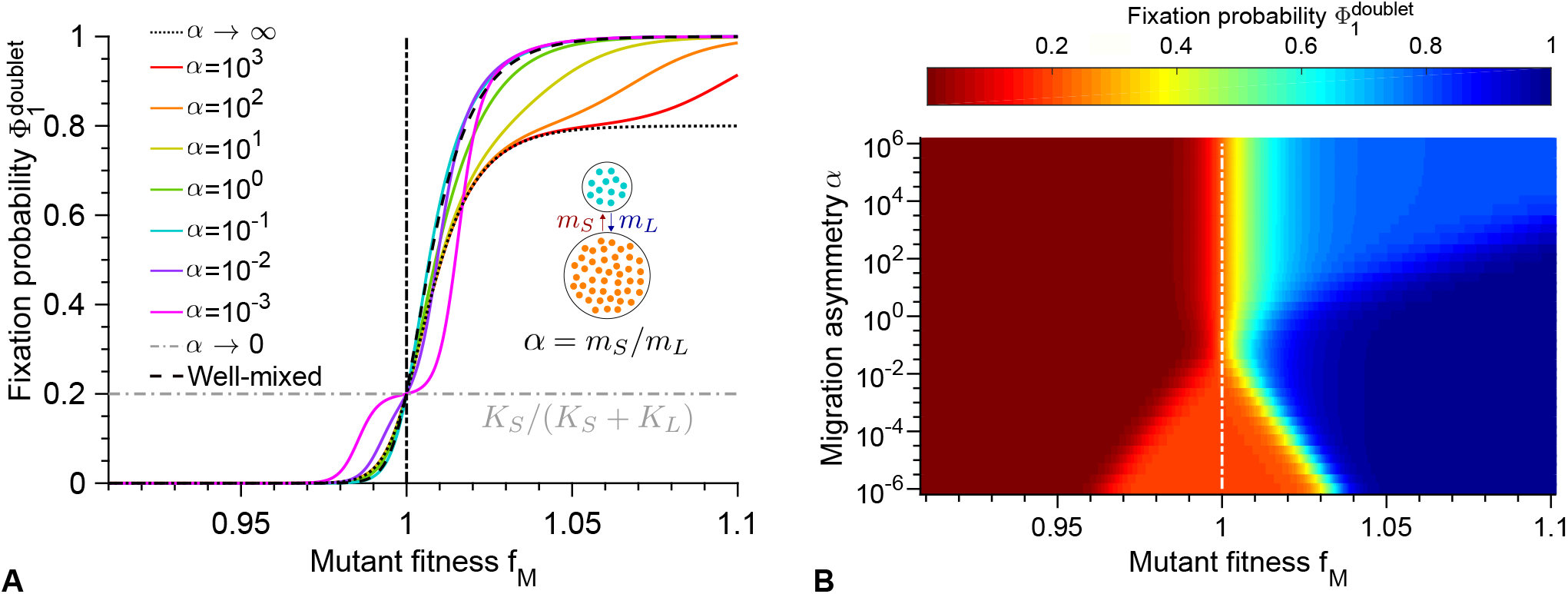
Fixation probability for the doublet. **A**: Fixation probability 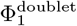 of mutants in a doublet versus mutant fitness *f*_*M*_, starting with one fully mutant deme chosen proportionally, with different migration rate asymmetries *α* = *m*_*S*_/*m*_*L*_, complementing those shown in Fig. S7. Data for the well-mixed population is shown as reference, with same total population size and initial number of mutants. Curves represent analytical predictions in Eqs. S84, S85 and S87. **B**: Heatmap of the same fixation probability versus mutant fitness *f*_*M*_ and migration rate asymmetry *α* = *m*_*S*_/*m*_*L*_. Parameter values in both panels: *K*_*S*_ = 100, *K*_*L*_ = 400 (hence *K*_*L*_ = (*D* − 1)*K*_*S*_ with *K*_*S*_ = *K* = 100 and *D* = 5), *f*_*W*_ = 1, *g*_*W*_ = *g*_*M*_ = 0.1. Vertical dash-dotted lines represent the neutral case *f*_*W*_ = *f*_*M*_.

### B. Fixation probability

#### 1. General expression

In order to calculate the fixation probability of the mutant type in the doublet, let us first consider the case where the small deme, whose carrying capacity is denoted by *K*_*S*_, is fully mutant, while the large deme, whose carrying capacity is denoted by *K*_*L*_, is fully wild-type. Recall that the migration rate per individual from the small deme to the large one is *m*_*S*_, and that from the large to the small deme by *m*_*L*_. We start from exactly one fully mutant deme. If an *M* individual migrates from the small deme to the large deme and fixes, then the mutant type fixes in the whole population. The probability that this occurs upon a given migration event reads

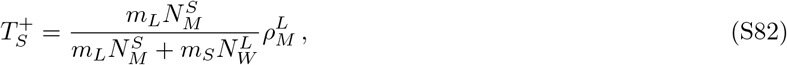

where 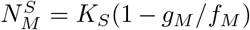 (respectively 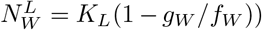 is the equilibrium size of the small mutant deme (respectively of the large wild-type deme) and 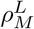 is the fixation probability of a mutant in the large wild-type deme, given by Eq. S7 with 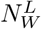 instead of *N*_*W*_. Similarly, if a *W* individual migrates to the small deme and fixes, then the wild-type fixes in the whole population. The probability that this occurs upon a given migration event reads

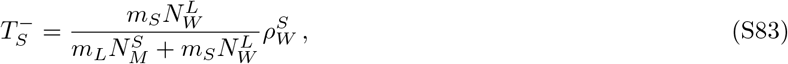

where 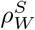 is the fixation probability of a wild-type individual in the small mutant deme, given by Eq. S10 with 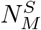 instead of *N*_*M*_. Then, the fixation probability of the mutant type, starting from a small mutant deme and a large wild-type deme, reads

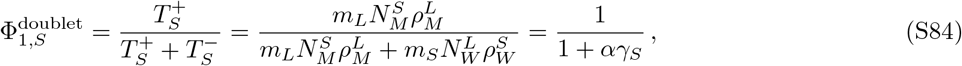

where *α* = *m*_*S*_/*m*_*L*_ and 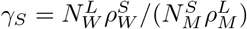.

Similarly, in the case where the structured population starts from a large mutant deme, while the small deme is wild-type, we get the fixation probability

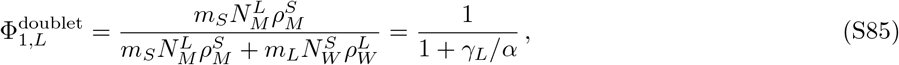

where 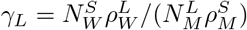.

Next, consider the case where one mutant individual starts in a deme with a probability proportional to the size of the deme, which corresponds to the realistic case of mutations happening randomly upon division. The fixation probability of such a single mutant reads:

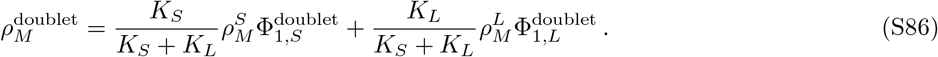

In the rest of this work, which focuses on structured populations made of demes of identical sizes, we consider the fixation probability Φ_1_ starting from one fully mutant deme, which then needs to be multiplied by *ρ*_*M*_ to obtain that of one mutant individual. Here, we will consider the analogous quantity

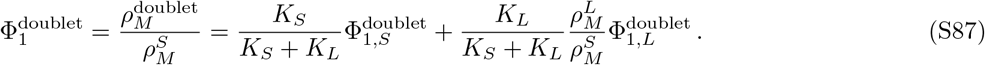

In the particular case where *K*_*L*_ = (*D* − 1)*K*_*S*_, so that the total carrying capacity of the subdivided population is *DK*_*S*_, denoting *K*_*S*_ by *K*, considering 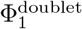 allows for a direct comparison to Φ_1_ in the other structures considered here, comprising *D* demes of carrying capacity *K*.

#### 2. Expansion for relatively small mutational effects

For the sake of simplicity, here we assume that *K*_*L*_ = (*D* − 1)*K*_*S*_, so that the total carrying capacity of the subdivided population is *DK*_*S*_, and we further denote *K*_*S*_ by *K*. Consider the regime where *ϵ* ≪ 1 and *N*_*W*_ |*ϵ*| + 1 but *N*_*W*_ *ϵ*^2^ ≪ 1. Then, if *ϵ* > 0, Eqs. S84, S85 and S87 yield

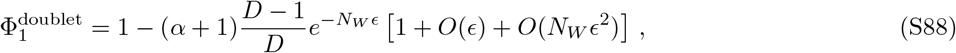

which gives, employing Eq. S19,

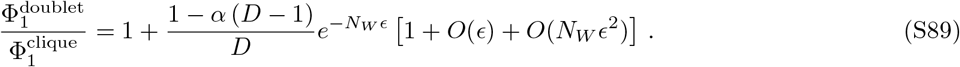

Thus, in this case, assuming *D* > 1, we have 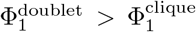 if *α* < 1/(*D* − 1), whereas 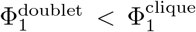 if *α* > 1/(*D* − 1). Now if *ϵ* < 0, Eqs. S84, S85 and S87 yield

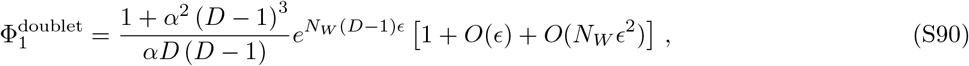

which gives, employing Eq. S20,

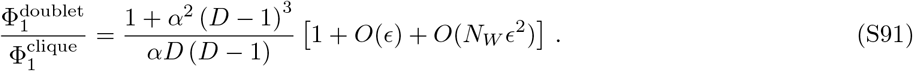

Then assuming *D* > 2, studying the function *G*:*α* → [1 + *α*_2_(*D* − 1)^3^]/[*α D*(*D* − 1)] demonstrates that 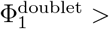 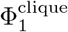 if *α* < 1/(*D* − 1)^2^ or *α* > 1/(*D* − 1), whereas 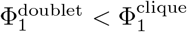 if 1/(*D* − 1)^2^ < *α* < 1/(*D* − 1). Therefore, in the regime where *ϵ* ≪ 1 and *N*_*W*_ *ϵ* ≫ 1 but *N*_*W*_ *ϵ*^2^ ≪ 1, the doublet is an amplifier of selection with respect to the clique for 1/(*D* − 1)2 < *α* < 1/(*D* − 1), and a suppressor of selection for *α* > 1/(*D* − 1). Finally, for *α* < 1/(*D* − 1)2, it behaves as a suppressor for *E* < 0 and as an amplifier for *ϵ* > 0.

#### 3. Expansion for extremely asymmetric migrations

If *α* → 0, then Eqs. S84, S85 and S87 yield

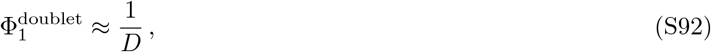

while if *α*→ ∞, they give

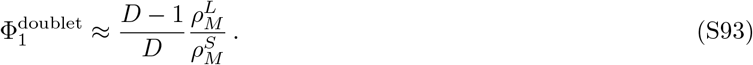

## VII. CONSTANT DEME SIZE APPROXIMATION

In our model, we consider that the number of individuals in each deme is not fixed, but there is a carrying capacity *K* per deme. In a deterministic description, valid for large populations, if there is only one type of individuals, the number *N* of individuals at time *t* follows the ordinary differential equation:

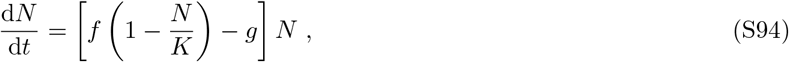

where *f* represents fitness, *g* death rate and *K* carrying capacity. If *f* > *g*, Eq. S94 yields a nonzero steady-state population size, namely *K*(1 − *g/f*). In a stochastic description, a finite-size microbial population with a logistic growth rate and a constant death rate fluctuates around the deterministic steady-state average population size *K*(1 − *g/f*) after a transient time depending on initial conditions and before eventually going extinct (after a very long time if it carrying capacity is not small) [56, 57]. Therefore, in our analytical studies, we often employ the steady-state population sizes of wild-type and mutant demes, denoted by *N*_*W*_ and *N*_*M*_ respectively:

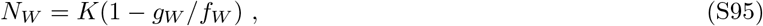

and

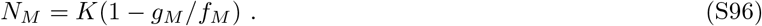

Furthermore, for simplicity, we approximate fixation probabilities in each deme by their values computed at fixed population size within the Moran process [26, 27]. The fixation probability of a single mutant (resp. wild-type) in a wild-type (resp. mutant) deme of steady-state size *N*_*W*_ (resp. *N*_*M*_) is then given by Eq. S7 (resp. Eq. S10). This approximation is expected to be reasonable for large enough steady-state deme sizes. This is confirmed by Fig. S9, where the constant-size approximation from Eq. S7 is compared to results from stochastic simulations of the evolutionary dynamics of a mutant in a population of *W* individuals with variable population size, and to a numerical resolution of the Master equation for variable population size, based on Ref. [58]. In the cases with variable population size, we use a carrying capacity *K* and a steady-state size *N*_*W*_ = *K*(1 − *g*_*W*_/*f*_*W*_), as in the rest of our work.

**FIG. S9.**
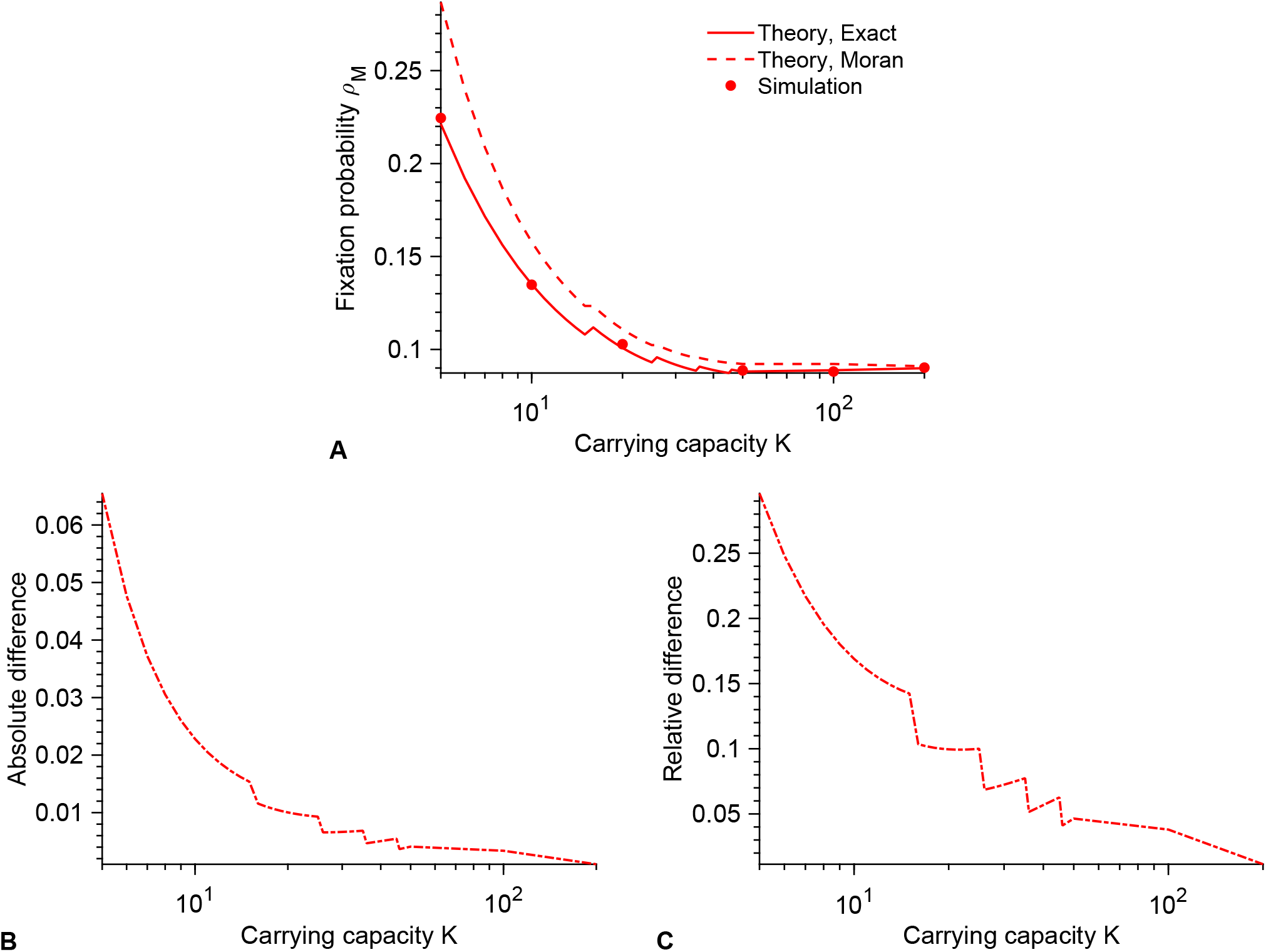
Constant deme size approximation. Fixation probability *ρM* of a mutant in a population of wild-type individuals with carrying capacity *K* and steady-state size *N*_*W*_ = *K*(1 − *g*_*W*_/*f*_*W*_). Markers: averages over 10^4^ stochastic simulations. Solid line: numerical resolution of the Master equation, see Eq. 3 in Ref. [58]. Dashed line: constant-size approximation employed in this work, in the framework of the Moran process [26, 27] (see Eq. S7). Parameter values: *f*_*W*_ = 1, *f*_*M*_ = 1.1, *g*_*W*_ = *g*_*M*_ = 0.1. Absolute and relative differences between the dashed and solid lines of panel **A** are shown in panels **B** and **C**. Discontinuities arise from the need to set the constant population size in the Moran to an integer value, while the steady-state size is not necessarily an integer – this is done by truncation.

## VIII. SIMULATION METHODS

Implementations of our simulations in the C programming language are freely available at https://doi.org/10.5281/zenodo.5126699.

Our numerical simulations are performed using a Gillespie algorithm that is exact and does not involve any artificial discretization of time [59, 60]. We focus on the regime where deme sizes *N*_*i*_ fluctuate weakly around their deterministic steady-state values, namely *N_i_ = K_i_*(1 − *g*_*a*_/*f*_*a*_) if all microbes in deme *i* are of type *a*. Thus, we start our simulations at these sizes, and we consider *K*_*i*_ large enough for stochastic extinctions not to occur within the timescales studied. In most cases, we start our simulations with one fully mutant deme, while all others are fully wild-type, because this describes the second step in the fixation of a mutant (after it has fixed in a deme) in the rare migration regime. Note however that our stochastic simulations are valid beyond the rare migration regime and allow us to test the validity of this assumption and to go beyond this regime. We consider a structured population of *D* demes labeled *i* = 1, 2, …, *D*, and denote by *N*__W,i__ and *N*_*M,i*_ the respective numbers of *W* and *M* individuals in deme *i*.

The elementary events that can happen are reproduction, death and migration of an individual of either type:

- 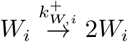 : Reproduction of a wild-type microbe in deme *i* with rate 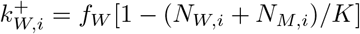.
- 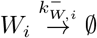: Death of a wild-type microbe in deme *i* with rate 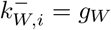.
- 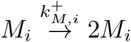: Reproduction of a mutant microbe in deme *i* with rate 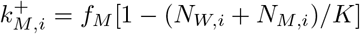.
- 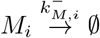 : Death of a mutant microbe in deme *i* with rate 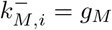. Note that we take *g*_*M*_ = *g*_*W*_ throughout.
- 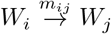: Migration of a wild-type microbe from deme *i* to deme *j* with rate *m*_*ij*_.
- 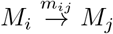 : Migration of a mutant microbe from deme *i* to deme *j* with rate *m*_*ij*_.

The total rate of events is given by 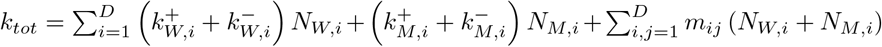.

Simulation steps are as follows:

1. Initialization: All of the *D* demes start from either *N*_*W*_ = *K*(1 − *g*_*W*_/*f*_*W*_) wild-type microbes or *N*_*M*_ = *K*(1 − *g*_*M*_/*f*_*M*_) mutant microbes, at time *t* = 0.
2. Monte Carlo step: Time *t* is incremented by Δ*t*, sampled from an exponential distribution with mean 1/*k*_*tot*_. The next event to occur is chosen proportionally to its probability *k/k*_*tot*_, where *k* is its rate, and is executed.
3. We go back to Step 2 unless only one type of individuals, either *W* or *M*, remains in the population, which corresponds to fixation of one type. Simulation is ended when fixation occurs.

## Notes

### Competing Interest Statement

The authors have declared no competing interest.

